# The immune receptor SNC1 monitors helper NLRs targeted by a bacterial effector

**DOI:** 10.1101/2023.03.23.533910

**Authors:** Ming-Yu Wang, Jun-Bin Chen, Rui Wu, Hai-Long Guo, Yan Chen, Zhen-Ju Li, Lu-Yang Wei, Chuang Liu, Sheng-Feng He, Mei-Da Du, Ya-long Guo, You-Liang Peng, Jonathan DG Jones, Detlef Weigel, Jian-Hua Huang, Wang-Sheng Zhu

**Author notes:** Correspondence (W.Z.) and (J.H.). Department of Plant & Environmental Studies, Copenhagen University, 1871 Frederiksberg, Denmark (Rui Wu). These authors contributed equally (Ming-Yu Wang, Jun-Bin Chen, Rui Wu, Hai-Long Guo).

## Abstract

Plants deploy intracellular receptors to counteract pathogen effectors that suppress cell-surface receptor-mediated immunity. To what extent pathogens manipulate also immunity mediated by intracellular receptors, and how plants tackle such manipulation, remains unknown. *Arabidopsis thaliana* encodes three very similar ADR1 class helper NLRs (ADR1, ADR1-L1 and ADR1-L2), which play key roles in plant immunity initiated by intracellular receptors. Here, we report that *Pseudomonas syringae* AvrPtoB, an effector with E3 ligase activity, can suppress ADR1-L1- and ADR1-L2-mediated cell death. ADR1, however, evades such suppression by diversification of two ubiquitination sites targeted by AvrPtoB. The intracellular sensor NLR SNC1 interacts with and guards the CC_R_ domains of ADR1-L1 and ADR-L2. Removal of ADR1-L1 and ADR1-L2 or delivery of AvrPtoB activates SNC1, which then signals through ADR1 to trigger immunity. Our work not only uncovers the long sought-after physiological function of SNC1 in pathogen defense, but also that reveals how plants can use dual strategies, sequence diversification and a multiple layered guard-guardee system, to counteract pathogen attack on core immunity functions.

## INTRODUCTION

Plants are constantly threatened by pathogens. To impede pathogen invasion, plants deploy plasma membrane-localized pattern-recognition receptors (PRRs) that initiate pattern-triggered immunity (PTI) upon detection of conserved molecular patterns diagnostic of pathogens. To enable successful invasion, pathogens in turn deliver effectors into plant cells to manipulate components of PTI. To antagonize the action of effectors, plants evolved intracellular nucleotide-binding domain leucine-rich repeat receptors (NLRs), which detect effectors or their effects on host proteins. The outcome is an enhanced immune response known as effector-triggered immunity (ETI). ETI usually culminates in programmed cell death called hypersensitive response (HR), a hallmark of ETI^1,2^. Recent studies have revealed at the molecular level how PTI and ETI are interlinked, with PTI and ETI potentiating each other^3–6^.

NLRs are classified into TIR-NLRs (TNLs), CC-NLRs (CNLs), and CC_R_-NLRs (RNLs), based on their N termini. RNLs are considered to function as helper NLRs downstream of sensor NLRs including most TNLs and some CNLs, which can directly or indirectly recognize effectors. Helper NLRs are encoded by three gene families, each with a different founding member: *ADR1* (*ACTIVATED DISEASE RESISTANCE 1*), *NRG1* (*N Requirement Gene 1*), and *NRC* (NLR Protein Required For Hypersensitive-response-associated Cell Death). *ADR1* homologs are ubiquitously present in angiosperm genomes, while the *NRG1* and *NRC* families are limited to dicots and Solanaceae, respectively^7^. The *Arabidopsis thaliana* genome encodes three unequally members of the ADR1 family: including ADR1, ADR1-L1 and ADR1-L2^7^. Like activated ZAR1 and Sr35 as well as NRG1, autoactive ADR1 can form Ca^2+^-permeable influx channels that activate cell death^8–10^. In addition, ADR1s form complexes with EDS1 (ENHANCED DISEASE SUSCEPTIBILITY 1)-PAD4 (PHYTOALEXIN DEFICIENT 4) heterodimers^3,11^. Similar to the *eds1* mutant, *adr1 adr1-L1 adr1-L2* triple mutants are highly susceptible to virulent *Pseudomonas syringae* as well as avirulent pathogens, resistance to which relies primarily on TNLs, but also some CNLs^12,13^. EDS1-PAD4-ADR1 complexes are also required for full PTI responses triggered by elicitor nlp20^3,6^. Taken together, these findings suggest that ADR1s play a key role in ETI and PTI.

*SNC1* (*SUPPRESSOR OF NPR1-1, CONSTITUTIVE 1*) encodes an extensively studied canonical sensor TNL^14^. Overexpression of wild-type *SNC1* activates salicylic acid (SA)-dependent defense responses^15^, and gain-of-function mutations in the coding sequence can suppress disease susceptibility of *npr1-1* mutants, which are defective in systemic acquired resistance (SAR)^14,16^. Subsequent studies on SNC1 uncovered complex control of NLRs, including epigenetic regulation, alternative splicing, intracellular trafficking, post-translational modification, and structural variation at SNC1 itself^17,18^. Inactivation of SNC1 restores elevated disease resistance seen in a range of autoimmune mutants with defects in very different types of genes^17^. Remarkably, even though SNC1 has become a powerful model to understand many different aspects of NLR regulation, its physiological roles in plant immunity, if any, have remained elusive.

An important role of pathogen effectors is to antagonize PTI components, with some Type III Secretion System (T3SS) effectors of *P. syringae* also suppressing ETI^19–21^. For example, HopI1 greatly dampens HR triggered by several other effectors by unknown mechanisms^21^. A recent reverse genetic screen identified five effectors from oomycetes and nematodes that suppress cell death triggered by NLRs Prf or Rpi-blb2 in *N. benthamiana*^22^.Among these effectors, SS15 exerts its effects by inhibiting the intramolecular rearrangements of NRC2, which prevents its oligomerization and activation^23^, while AVRcap1b dampens NRC2 and NRC3 function through the membrane trafficking-associated protein NbTOL9a (Target of Myb 1-like protein 9a)^22^. From these studies it is clear that much is still to be learned about how pathogens suppress ETI and how plants in turn counteract such suppression.

Here, we report that the *P. syringae* effector AvrPtoB, an E3 ligase induces SNC1 oligomerization by ubiquitinating and promoting degradation of the *A. thaliana* helper NLR ADR1-L1. Two non-synonymous substitutions in the CC_R_ domain allow the ADR1-L1 homolog ADR1 to evade AvrPtoB-mediated ubiquitination. ADR1-L1 itself is guarded by the sensor NLR SNC1. The autoimmunity of *adr1-L1-1* single and *adr1-L1-1 adr1-L2* double mutants is suppressed by inactivation of *ADR1*, indicating that ADR1 acts downstream of ADR1-L1 and ADR1-L2. Together, we demonstrate that the sensor NLR SNC1 recognises AvrPtoB by guarding ADR1-L1 and ADR1-L2, then signals through ADR1 for immune responses. Our findings uncover a plant mechanism for counteracting ETI suppression by bacterial effectors, illustrating yet another layer of plants neutralizing pathogen effectors.

## RESULTS

### AvrPtoB induces ADR1-L1 protein degradation

We use *Pseudomonas syringae* (*Pst*) pv. tomato DC3000 model to study the interaction between the plant immune system and pathogen effectors. To identify *Pst* DC3000 effectors that suppress the activity of the essential ETI component ADR1-L1 from *A. thaliana* (hereafter Arabidopsis), we first generated an autoactive ADR1-L1 variant (ADR1-L1^D489V^), which triggers robust cell death in *N. benthamiana* **(Extended Data Fig. 1a)**. We co-expressed this variant, with a mutation in the MHD regulatory motif, in individual combinations with 31 of the 36 *Pst* DC3000 effectors in *N. benthamiana* in search for effectors that might dampen ADR1-L1^D489V^-triggered cell death **(Fig. 1a)**. Only AvrPtoB did so completely **(Fig. 1b, Extended Data Fig. 1b)**.

**Fig. 1.**
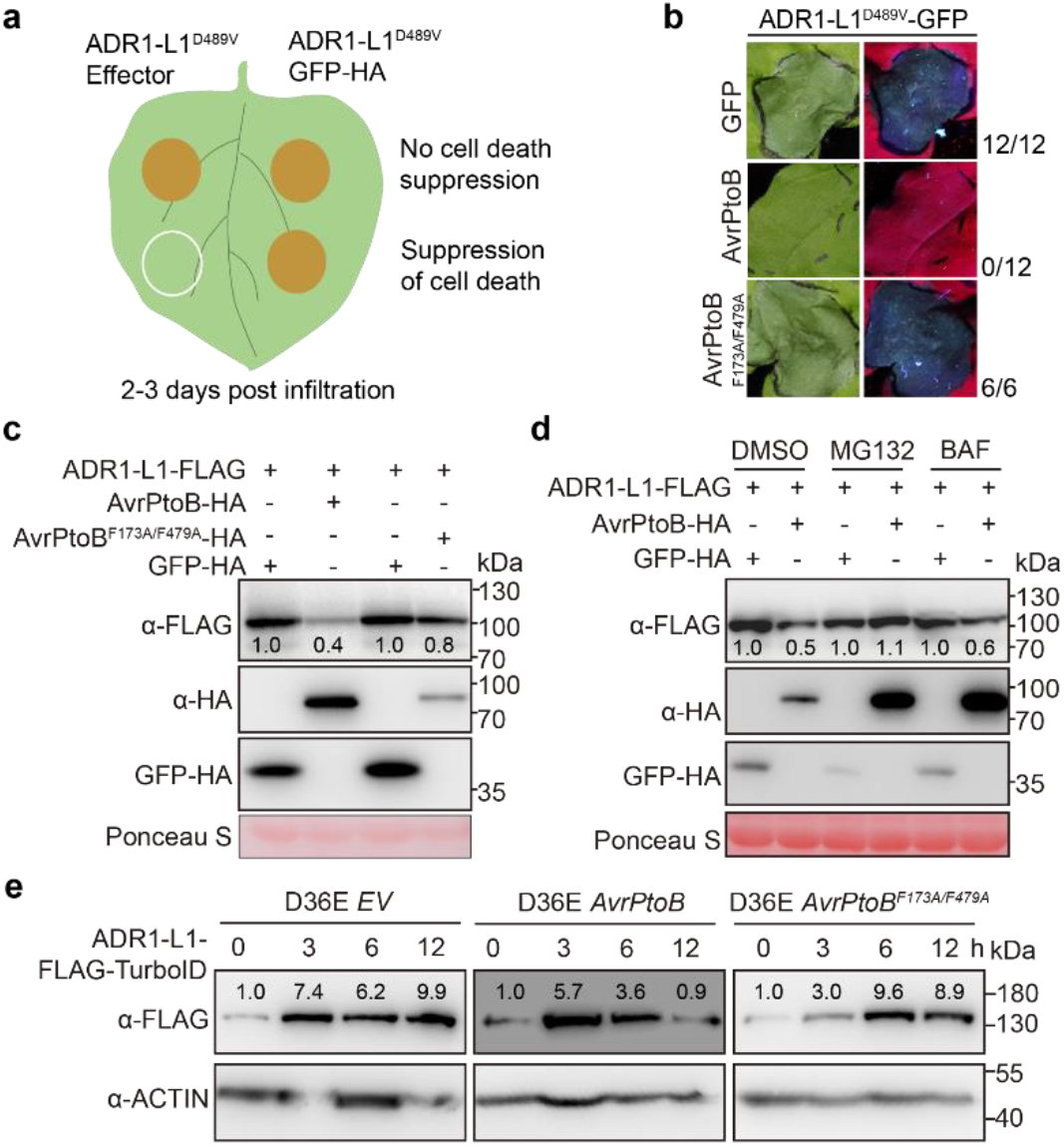
AvrPtoB suppresses ADR1-L1-triggered HR and induces the degradation of ADR1-L1. **a**, Schematic diagram of the screen of *Pst* DC3000 effectors that suppress HR triggered by transient expression of ADR1-L1^D489V^ in *N. benthamiana*. **b**, E3 ligase activity of AvrPtoB is required for suppression of HR triggered by ADR1-L1^D489V^. Numbers on the far right indicate leaves showing obvious HR over all infiltrated leaves. **c**, E3 ligase activity is required for AvrPtoB inducing degradation of ADR1-L1. **d**, The 26S proteasome inhibitor MG132 blocks degradation of ADR1-L1 induced by AvrPtoB. **e**, AvrPtoB induces degradation of ADR1-L1-FLAG-TurboID in four-week-old transgenic Arabidopsis plants. Numbers indicate arbitrary densitometry units of corresponding bands after normalization to the left-most ADR1-L1-FLAG-TurboID band of each immunoblot. Experiments were performed three times, with similar results.

AvrPtoB is a U-box E3 ligase^24^. The E3 ligase-dead variant AvrPtoB^F173A/F479A^ (ref. <u>24</u>) did not suppress ADR1-L1^D489V^ -triggered cell death **(Fig. 1b)**, indicating that AvrPtoB uses its E3 ligase activity to manipulate ADR1-L1 function. Levels of ADR1-L1-FLAG protein in *N. benthamiana* leaves were substantially reduced when co-expressed with AvrPtoB-HA, but not when co-expressed with the catalytically inactive AvrPtoB^F173A/F479A^ variant **(Fig. 1c)**. Such reduction was alleviated in the presence of the 26S proteasome inhibitor MG132, but not in the presence of BAF, which inhibits protein degradation by the autophagy pathway **(Fig. 1d)**. These results suggest that AvrPtoB triggers ADR1-L1 degradation in an E3 ligase activity-dependent manner via the 26S proteasome pathway.

To further confirm the degradation of ADR1-L1 catalysed by AvrPtoB, wild-type and catalytically inactive variants were delivered by the effectorless *Pst* DC3000D 36E strain^19^ into Arabidopsis ADR1-L1-FLAG-TurboID plants. ADR1-L1-FLAG-TurboID protein levels had increased at 3 hours post infiltration (hpi) for all treatments **(Fig. 1e)**, likely due to activation of PTI by *Pst* DC3000 D36E. ADR1-L1-FLAG-TurboID protein level had levelled off at 6 hpi when plants were infiltrated with *Pst* DC3000 D36E expressing AvrPtoB, and decreasing further at 12 hpi **(Fig. 1e)**. In contrast, no changes in ADR1-L1-FLAG-TurboID protein level were observed at 6 and 12 hpi when plants were infiltrated with *Pst* DC3000 D36E expressing AvrPtoB^F173A/F479A^ **(Fig. 1e)**. Taken together, these observations suggest that AvrPtoB induces the degradation of ADR1-L1 in Arabidopsis during pathogen infection.

### The CC_R_ domain determines AvrPtoB targeting

Since the three ADR1 members share similar functions in regulating intracellular receptor-dependent immune responses, we wondered whether AvrPtoB also compromised the stability of ADR1 and ADR1-L2 as well as the ability of their autoactive variants to trigger HR. In contrast to ADR1-L1^D489V^, HR triggered by ADR1^D461V^ was only rarely suppressed, and HR triggered by ADR1-L2^D484V^ was only slightly suppressed by AvrPtoB **(Fig. 2a)**, even though the co-immunoprecipitation (Co-IP) and split-luciferase complementation (SLC) had indicated that AvrPtoB can interact with all ADR1 members **(Fig. 2b, Extended Data Fig 2a, b)**. In agreement, ADR1 protein levels in *N. benthamiana* were not affected by AvrPtoB **(Extended Data Fig 2c)**. The weak effects on ADR1-L2 protein abundance may be due to the mild suppression of ADR1-L2 by AvrPtoB, which is consistent with the modest impairment of ADR1-L2^D484V^-mediated cell death by AvrPtoB **(Fig. 2a, Extended Data Fig. 2c)**. Furthermore, infiltration of *Pst* DC3000 D36E carrying AvrPtoB did not alter the protein level of either ADR1-FLAG-TurboID or ADR1-L2-Flag-TurboID in Arabidopsis **(Extended Data Fig. 2d)**. These results suggest that AvrPtoB affects the stability of ADR1 family members as well as the HR they trigger in a homolog-specific manner.

**Fig. 2.**
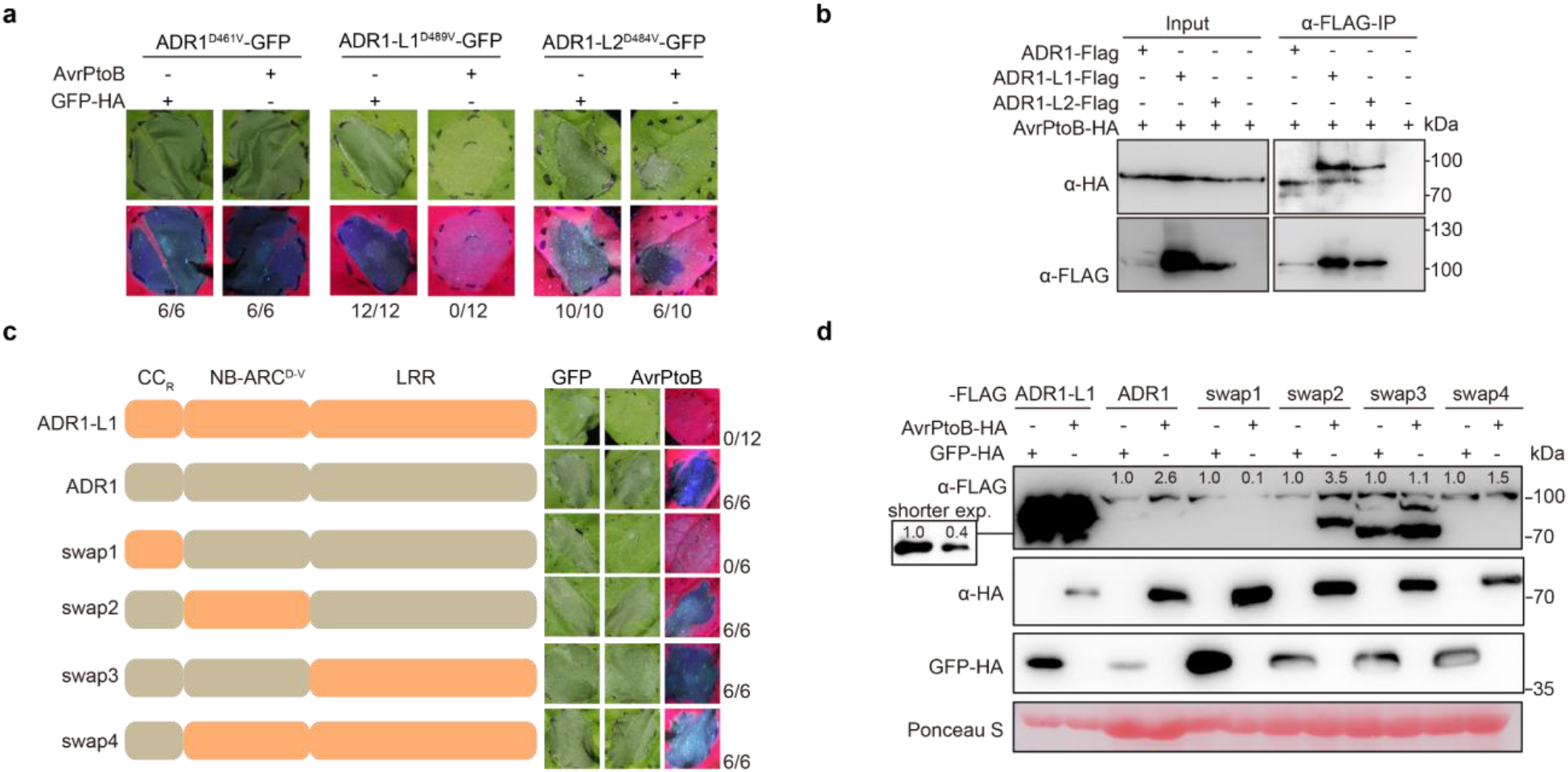
The CC_R_ domains are responsible for the differential AvrPtoB suppression of ADR1 and ADR1-L1 activity. **a**, AvrPtoB differentially suppresses HR triggered by autoactivate ADR1-L1 homologs. ADR1^D461V^, ADR1-L1^D489V^, and ADR1-L2^D484V^ were transiently co-expressed with GFP-HA and AvrPtoB-HA in *N. benthamiana*. **b**, AvrPtoB associates with the three ADR1 homologs, as shown by Co-IP in *N. benthamiana*. **c**, Domain swapping shows that the CC_R_ domains of ADR1 homologs determine susceptibility to AvrPtoB suppression. **d**, Swapping the CC_R_ domains between ADR1 and ADR1-L1 switches the AvrPtoB-susceptibility of ADR1 and ADR1-L1. Numbers on the bottom **(a)** or far right **(c)** indicate leaves with HR over all infiltrated leaves. Experiments were performed three times, with similar results.

To identify the causal domains responsible for differential suppression of ADR1- and ADR1-L1-triggered HR by AvrPtoB, we swapped the CC_R_, NB-ARC, and LRR domains between ADR1-L1^D489V^ and ADR1^D461V^. Interchange of the CC_R_ domain, but not the NB-ARC and LRR domains, made ADR1^D461V^-triggered cell death responsive to AvrPtoB, and at the same time made ADR1-L1^D489V^-triggered cell death insensitive to AvrPtoB **(Fig. 2c)**. In agreement, ADR1^D461V^ with the CC _R_^ADR1-L1^ domain, but not with the NB-ARC^ADR1-L1^ or LRR^ADR1-L1^ domains, accumulated to a lower level in the presence of AvrPtoB, while the levels of ADR1-L1^D489V^ with the CC_R_ ^ADR1^ domain were insensitive to the presence of AvrPtoB **(Fig. 2d)**. These results indicate that the CC_R_ domain determines the specificity of AvrPtoB-mediated suppression of ADR1-L1 activity.

As sequence differences in the CC_R_ domains are responsible for differential effects of AvrPtoB on ADR1 homologs, we tested whether AvrPtoB can inhibit also the cell death caused by transient expression of only the CC_R_ domain of ADR1 homologs in *N. benthamiana*^10,25^. Similar to AvrPtoB effects on the autoactive full-length variants, AvrPtoB did not affect CC_R_^ADR1^-triggered cell death, slightly suppressed CC_R_^ADR1-L2^-triggered cell death, and abolished CC _R_^ADR1-L1^-triggered cell death **(Fig. 3a)**. This was paralleled by AvrPtoB having little impact on the protein levels of CC_R_ ^ADR1^ and CC_R_ ^ADR1-L2^, but causing a substantial reduction of CC_R_ ^ADR1-L1^ levels **(Extended Data Fig. 3a)**. Thus, the effects of AvrPtoB on both protein accumulation and cell death-inducing ability are similar between the CC_R_ domains and full-length ADR1 homologs **(Fig. 2b, 3a, Extended Data Fig. 2d, 3a)**.

**Fig. 3.**
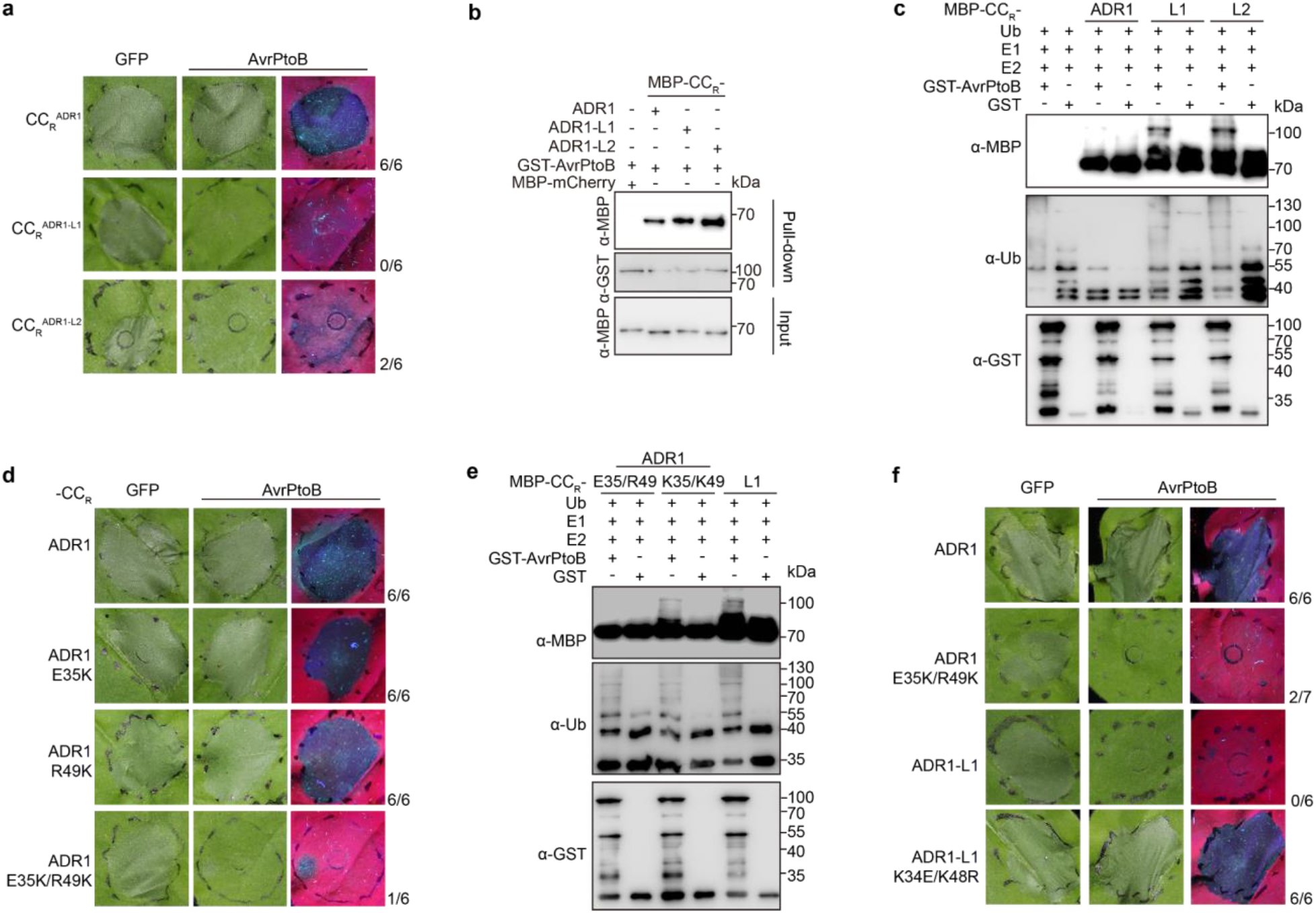
Two lysine residues in the CC_R_ domain are required for AvrPtoB-dependent suppression of ADR1-L1^D489V^ activity. **a**, AvrPtoB fully and partially suppresses HR triggered by CC^RADR1-L1^ and CC^RADR1-L2^, but not at all HR triggered by CC_RADR1_ in *N. benthamiana*. **b**, AvrPtoB associates with the CC_R_ domains of the three ADR1 homologs *in vitro*, as shown by pull-down assays with proteins purified from *E. coli*. **c**, AvrPtoB ubiquitinates CC_RADR1-L1_ and CC_RADR1-L2_, but not CC_RADR1_, as shown by *in vitro* ubiquitination assay with proteins purified from *E. coli*. **d**, AvrPtoB suppresses HR triggered by the E35K/R49K mutations in *N. benthamiana*. **e**, AvrPtoB ubiquitinates CC_RADR1_ with E35K/R49K but not wild-type CC_RADR1_, as shown by *in vitro* ubiquitination with proteins purified from *E. coli*. **f**, AvrPtoB suppresses HR triggered by full-length ADR1^D461V^ with E35K/R49K mutations in *N. benthamiana*. Conversely, AvrPtoB no longer suppresses HR triggered by ADR1-L1^D489V^ upon introduction of the K34E/K48R mutations. Numbers on the right **(a, d**, and **f)** indicate leaves with HR over all infiltrated leaves tested. Experiments were performed three times, with similar results.

Because the MBP-tagged CC_R_ domains of all three ADR1 homologs were similarly pulled down by purified AvrPtoB-GST, interaction of AvrPtoB with CC_R_ domains **(Fig. 3b)** is apparently not sufficient for AvrPtoB to promote protein degradation **(Extended Data Fig. 3a)**, likely due to differential ubiquitination of CC_R_ ^ADR1^ and CC_R_ ^ADR1-L1^ by AvrPtoB. An in vitro assay confirmed that AvrPtoB can efficiently ubiquitinate CC_R_ ^ADR1-L1^ and CC_R_ ^ADR1-L2^ but not CC _R_^ADR1^ **(Fig. 3c)**. This is consistent with AvrPtoB being able to at least partially suppress cell death triggered by CC _R_^ADR1-L1^ and CC_R_^ADR1-L2^, and CC _R_^ADR1^ being immune to AvrPtoB. Our results indicate that CC _R_^ADR1^escapes suppression of AvrPtoB by evading AvrPtoB-catalysed ubiquitination.

To identify the residues that allow CC_R_ ^ADR1^ to avoid becoming ubiquitinated, we generated chimeric CC_R_ proteins by swapping the first 50 amino acids between CC_R_ ^ADR1-L1^ and CC _R_^ADR1^, then co-expressed the chimeric CC_R_ proteins with AvrPtoB in *N. benthamiana* **(Extended Data Fig. 3b, c)**. While AvrPtoB failed to suppress cell death triggered by wild-type CC_R_^ADR1^, it abolished the cell death caused by the CC_R_ ^ADR1^ chimera with the first 50 amino acids of CC _R_^ADR1-L1^ **(Extended Data Fig. 3d)**.

Canonical ubiquitination occurs on lysine residues. The first 50 amino acids of ADR1-L1 contain only two lysines, K34 and K48, that are conserved in ADR1-L2. The CC_R_ domain from ADR1 instead features a glutamate (E35) and an arginine (R49) in these two positions **(Extended Data Fig. 3b)**. The E35 and R49 residues may enable ADR1 to evade being targeted by AvrPtoB. To test this hypothesis, we mutated E35 and R49 of the CC_R_^ADR1^ to lysine (E35K and R49K) and examined the effects of the two mutations on AvrPtoB susceptibility. When both E35K and R49K were introduced, cell death triggered by CC _R_^ADR1^ was dramatically inhibited by AvrPtoB **(Fig. 3d, Extended Data Fig. 3d)**. As expected, CC_R_^ADR1^ with E35K/R49K substitutions was ubiquitinated by AvrPtoB **(Fig. 3e)**. We also introduced these changes in the context of the full-length ADR1^D461V^ gain-of-function variant, which became susceptible to suppression by AvrPtoB as well **(Fig. 3f, Extended Data Fig. 3e)**. Conversely, when K34 and K48 of ADR1-L1^D489V^ were mutated to glutamate and arginine, ADR1-L1^D489V^-triggered cell death could no longer be suppressed by AvrPtoB **(Fig. 3f, Extended Data Fig. 3e)**. Taken together, our results indicate that the K34 and K48 residues are the functionally relevant sites in the CC_R_ domain of ADR1-L1 that are ubiquitinated by AvrPtoB. Because ADR1 features different residues in these positions, E35 and R49, it evades suppression of its activity by AvrPtoB.

To understand the evolutionary history of changes at the CC_R_ residues crucial for targeting by AvrPtoB, we reconstructed the phylogeny of 552 ADR1 homologs from angiosperms. The 117 Brassicaceae homologs form a single clade, indicating that diversification occurred only in the Brassicaceae, with the ADR1 clade apparently being younger than the ADR-L1 clade **(Extended Data Fig. 3f)**. Focusing on the two lysine residues targeted by AvrPtoB, we find that an ADR1/ADR1-L1/ADR1-L2 homolog from the sister lineage of Brassicaceae *Tarenaya hassleriana* at the base of the Brassicales encodes a lysine corresponding to position 48 in ADR1-L1, but not at position 34. In the Brassicaceae, the ADR-L1 and ADR-L2 homologs show similar profiles, with lysine being the most common residue at position 46/48, while lysine is found in that position only in a minority of ADR1 homologs. At position 32/34, several ADR1-L1/L2 homologs have a lysine, but lysine is never found at that position in ADR1 **(Extended Data Fig. 3g)**. Notably, lysines at these two positions are exceedingly rare in ADR1 homologs outside of the Brassicaceae, suggesting an unknown trade-off that led to the evolution of lysines at these positions in the Brassicaceae, despite these residues being targets of AvrPtoB.

### *adr1-L1* null mutants express constitutive immunity

The *adr1-L1-1* mutant, reported to carry a T-DNA insertion disrupting the first exon of *ADR1-L1*, was used in previous studies to characterize the effects of *ADR1-L1* knockout on plant immunity, with the conclusion that the mutant on its own has no major phenotypes^26,27^, although *adr1-L1-1* as well as two EMS-induced point mutations in *ADR1-L1, muse15-1* and *muse15-2*, enhance *snc1* gain-of-function autoimmune defects^27^. To confirm that *adr1-L1-1* is a knockout allele, we used an amplicon that spans the first and second exon of *ADR1-L1* to quantify mRNA expression in RT-qPCR assays. We found that the T-DNA mutant still expressed about 30% of the amount of *ADR1-L1* mRNA observed in wild type **(Extended Data Fig. 4a, b)**, indicating that *adr1-L1-1* is only a knockdown allele.

**Fig. 4.**
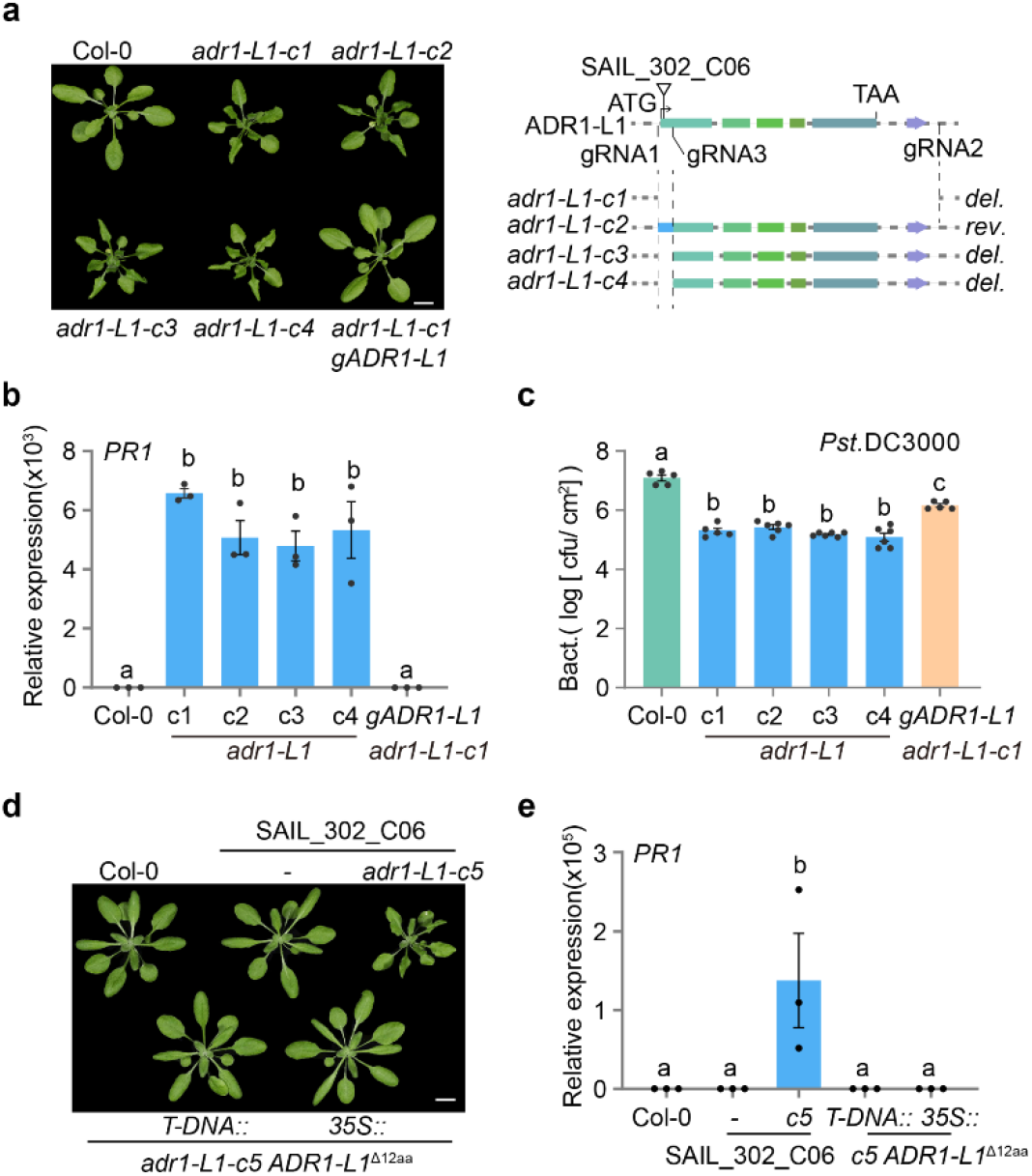
Inactivation of *ADR1-L1* causes autoimmunity. **a**, Left, four independent *adr1-L1* null mutants generated by CRISPR/Cas9 have typical autoimmune phenotypes, which are rescued by a genomic *ADR1-L1* copy (“*gADR1-L1*”). Right: diagram of T-DNA insertion in *adr1-L1-1*, the region targeted by guideRNAs (gRNAs) for CRISPR/Cas9-mediated inactivation, and the resultant *adr1-L1* null alleles. Scale bar: 10 mm. **b**, *PR1* expression is increased in *adr1-L1* mutants. *PR1* expression in plants in **(a)** was quantified by RT-qPCR. **c**, *adr1-L1* mutants have enhanced resistance to *Pst* DC3000 infection. **d**, The T-DNA mutant line *SAIL_302_C06* is a partial loss-of-function allele of *ADR1-L1*. Four-week-old plants are shown. Scale bar: 10 mm. **e**, *PR1* expression is also increased in the *adr1-L1-c5* mutant generated in the *adr1-L1-1* background. *PR1* expression in plants shown in **(d)** was quantified by RT-qPCR assays. Data in **(b, c, e)** represent the mean and standard error (n = 3, 5, and 3 biologically independent samples for (**b**), (**c**), and (**e**), respectively. *p* < 0.05, one-way ANOVA followed by Tukey’s post hoc test, letters indicate significantly different groups).

We generated a null mutant of *ADR1-L1, adr1-L1-c1*, by deleting the full coding region of *ADR1-L1* through CRISPR/Cas9 gene editing **(Fig. 4a)**. No *ADR1-L1* expression was detected in the mutant by RT-qPCR **(Extended Data Fig. 4c)**. Surprisingly, the *adr1-L1-c1* mutant was stunted and had curly leaves **(Fig. 4a)**, two hallmarks of autoimmunity in Arabidopsis^28^. To exclude the possibility that the phenotypes of *adr1-L1-c1* mutant were due to off-target effects of the CRISPR/Cas9 system, we transformed *ADR1-L1* driven by its native promoter into *adr1-L1-c1* mutants. Dwarfing and leaf curling were rescued in the *adr1-L1-c1* complementation lines **(Fig. 4a)**, confirming that the observed phenotypes are due to knockout of *ADR1-L1*. Three additional independent *adr1-L1* CRISPR/Cas9 mutants (*adr1-L1-c2, adr1-L1-c3, adr1-L1-c4*), which had either a small inversion or small deletions in the region encoding the CC_R_ domain, were also stunted in size and had curly leaves, mimicking the *adr1-L1-c1* mutants **(Fig. 4a)**.

We next quantified expression of the defense marker gene *PR1* to determine whether the phenotypes of the new *adr1-L1* mutants were indeed due to autoimmunity. *PR1* expression was increased in all four new *adr1-L1* mutants **(Fig. 4b)**, and this increase was reversed in the *adr1-L1-c1* complementation lines. In accordance, growth of the bacterial pathogen *Pst* DC3000 was impaired in the four new *adr1-L1* mutants, and this mutant phenotype was again rescued in the *adr1-L1-c1* complementation lines **(Fig. 4c)**. To confirm that the absence of reported phenotypes for the previously reported T-DNA allele^26,27^ did not result from differences in growth conditions, we grew it alongside the new *adr1-L1-c1* mutant, confirming that only the T-DNA knockdown allele appeared normal **(Extended Data Fig. 4d)**. Collectively, these results demonstrate that a complete knock out of *ADR1-L1* leads to spontaneous activation of immune signaling.

To investigate further why the T-DNA insertion in *adr1-L1-1* T-DNA causes only partial loss of function, we carried out further RT-PCR analyses, which showed that this allele produces a 5’ truncated transcript, with the T-DNA fragment providing a new start codon that should produce a nearly-full-length protein lacking only amino acids 2 to 13 (ADR1-L1^Δ12aa^) **(Extended Data Fig. 4b-f)**. Deleting *ADR1-L1* including the inserted T-DNA using CRISPR/Cas9 led to dwarfism and elevated *PR1* expression, which was rescued when the plants were transformed with a construct containing *ADR1-L1*^*Δ12aa*^ driven by the 3’ region of the T-DNA or the CaMV35S promoter **(Fig. 4d, e)**. These results confirm that *adr1-L1-1* is only a partial loss-of-function allele that does not cause autoimmunity.

### *adr1-L1* null mutant defects are *SNC1*-dependent

The defense marker *PR1*, which is greatly increased in *adr1-L1* null mutants, is regulated by salicylic acid (SA), and SA signaling in turn is protected by *EDS1* and *PAD4*^29^. To begin to uncover the mechanism underlying the spontaneous activation of immunity in *adr1-L1* null mutants, we first crossed *adr1-L1-c1* mutants to plants deficient for the salicylic acid biosynthesis gene *SID2* (*SALICYLIC ACID INDUCTION DEFICIENT 2*) or for *PAD4* and *EDS1*. The morphological defects of *adr1-L1-c1* were partially suppressed by *sid2-2* and fully suppressed by *eds1-2* and *pad4-1* **(Extended Data Fig. 5a)**.

**Fig. 5.**
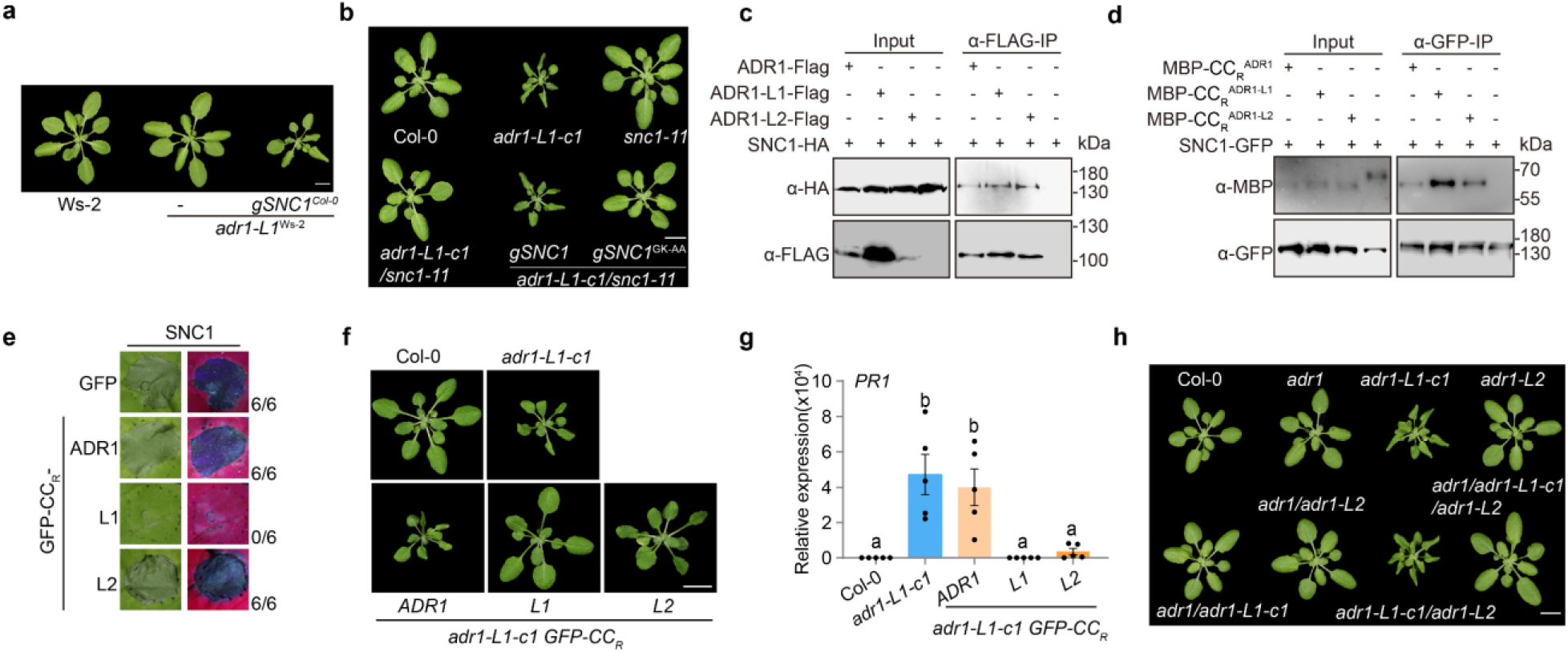
SNC1 guards ADR1-L1 and ADR1-L2 and signals through ADR1. **a**, The natural loss-of-function *SNC1* allele in Ws-2 suppresses growth defects of *adr1-L1* null mutants in Ws-2. Four-week-old plants of Ws-2, *adr1-L1*^*Ws-2*^ and *adr1-L1*^*Ws-2*^ transgenic line carrying an *SNC1* genomic fragment from Col-0. Scale bar: 10 mm. **b**, The loss-of-function *snc1-11* allele suppresses growth defects of the *adr1-L1*-*c1* null mutant in Col-0. This effect is reversed when a wild-type *SNC1* genomic fragment is introduced, but not the mutant *SNC1*^*GK-AA*^ variant. Scale bar: 10 mm. **c**, SNC1 associates with the three ADR1 homologs, as shown by Co-IP assays in *N. benthamiana*. **d**, SNC1 interacts with the CC_R_ domains of the three ADR1 homologs, as shown by semi-*in vitro* pull-down assays. SNC1-GFP and MBP-CC_R_ proteins were purified from *N. benthamiana* and *E. coli*, respectively. **e**, The CC_R_ domain of GFP-tagged ADR1-L1 efficiently suppresses SNC1-triggered HR in *N. benthamiana*. Numbers on the right indicate leaves with HR over all infiltrated leaves tested. **f**, Expression of GFP-tagged CC_R_ domains of ADR1-L1 and ADR1-L2 but not ADR1 suppress the growth defects of *adr1-L1-c1*. Representative four-week-old Arabidopsis T_1_ transgenic plants with *p35S::GFP-CC*_*R*^*ADR1*^_, *p35S::GFP-CC*_*RADR1-L1*_ *and p35S::GFP-CC*_*R*^*ADR1-L2*^_ in *adr1-L1-c1*, grown in 23°C. Scale bar, 10 mm. **g**, *PR1* expression of three-week old T_1_ transformants shown in **(f)**. Data represent the mean and standard error of five independent T_1_ transformants (n = 5 biologically independent samples, *p*<0.05, one-way ANOVA followed by Tukey’s post hoc test; letters indicate significantly different groups). **h**, Three-week-old *adr1-L1-c1* single and multiple mutants, grown at 23°C. Scale bar, 10 mm. Experiments in **(c-e)** were performed three times, with similar results

Because autoimmunity often results from inappropriate activation of NLR activity, we speculated that the autoimmune phenotype of *adr1-L1* mutants might result from genetic interaction with other NLRs. To identify such NLR candidates, we exploited the extensive variation in NLR complements in different Arabidopsis accessions^30^, and deleted *ADR1-L1* in the Arabidopsis accessions Est-1, C24 and Ws-2. Different from Col-0 and C24, inactivation of *ADR1-L1* in Ws-2 and Est-1 did not cause obvious morphological defects **(Fig. 5a)**. An F_2_ mapping population was generated by crossing *adr1-L1* (Ws-2) and *adr1-L1-c1* (Col-0). Genetic linkage analysis identified a single large-effect locus on chromosome 4 that suppressed *adr1-L1* autoimmune defects. Fine mapping narrowed the interval to a ∼130 kb region from 9.47 Mb to 9.60 Mb on chromosome 4 **(Extended Data Fig. 5b)**, which encompasses the *RPP4* cluster of *TNL* genes.

The *RPP4* cluster includes the intensively studied TNL gene *SNC1*, which is functional in Col-0, but not in Ws-2^31^, one of the two accessions in which the *adr1-L1* knockout phenotype is suppressed. To test whether *SNC1* is a natural modifier of *adr1-L1*, we transformed the *SNC1* (Col-0) genomic fragment into the *adr1-L1* (Ws-2) mutant. The transgenic plants resembled the *adr1-L1-c1* mutant of the Col-0 accession **(Fig. 5a)**. Furthermore, in Col-0, the *snc1-11* knockout allele suppressed morphological and molecular defects of *adr1-L1-c1* mutants **(Fig. 5b, Extended Data Fig. 5c-d)**, confirming that *SNC1* is the natural modifier of *ADR1-L1*. Dwarfism of the *adr1-L1-c1/snc1-11* mutant was restored by introducing the wild-type *SNC1* genomic fragment but not its P-loop mutant *SNC1*^*GK-AA*^ **(Fig. 5b)**. These results together showed that the *adr1-L1-c1* mutant defects are mediated by *SNC1*, most likely through activation of *SNC1* signaling.

### SNC1 guards ADR1-L1/L2 and signals through ADR1

The genetic interaction of *SNC1* and *ADR1-L1* prompted us to test their physical interaction. SNC1 was pulled down by all three ADR1 homologs in Co-IP assays in *N. benthamiana* **(Fig. 5c)**. In vitro pull-down experiments pointed to SNC1 interacting, likely with different affinities, with the CC_R_ domains of the three ADR1 homologs **(Fig. 5d)**.

Given the genetic and physical interaction between ADR1-L1 and SNC1, we hypothesized that SNC1, a sensor NLR, may guard ADR1-L1 through binding its CC_R_ domain, with loss of ADR1-L1 leading to SNC1 activation, as seen with some other NLRs that directly guard cellular targets^17^. Transient expression of SNC1 on its own triggered cell death in *N. benthamiana*, which could be suppressed by co-expression of GFP-CC_R_ ^ADR1-L1^ but not GFP-CC_R_ ^ADR1^ and GFP-CC_R_ ^ADR1-L2^ **(Fig. 5e, Extended Data Fig. 5f)**. In Arabidopsis, overexpression of *GFP-CC*_*RADR1-L1*_ completely suppressed the phenotypes of *adr1-L1-c1* mutants (T_1_ plants, n = 26). Overexpression of *GFP-CC*_*R*_^*ADR1-L2*^ could sometimes partially suppress *adr1-Ll-c1* phenotypes (7/28 T_1_ plants), while *GFP-CC*_*R*_^*ADR1*^ was ineffective (n = 56) **(Fig. 5f, g)**. We conclude that through monitoring the presence of their CC_R_ domains, SNC1 mainly guards ADR1-L1 and, to a lesser extent, ADR1-L2 but not ADR1. A minor role of SNC1 in guarding ADR1-L2 was further supported by the observation that the *adr1-L2* mutation slightly enhanced the *adr1-L1-c1* phenotype **(Fig. 5h, Extended Data Fig. 5h)**..

Phenotypic abnormalities in the *adr1-L1-c1* single and the *adr1-L1-c1/adr1-L2* double mutants were completely suppressed in the presence of the *adr1* mutation **(Fig. 5h)**. Taken together, these results indicate that ADR1-L1 and ADR1-L2 are guardees of SNC1, which signals via ADR1 to activate downstream responses.

### SNC1 recognises AvrPtoB through ADR1-L1

Structural studies^9,32–35^ have revealed how oligomerization of TNL proteins ROQ1 and RPP1, as well as CNL proteins ZAR1 and Sr35 is associated with their activation. We therefore used BN-PAGE to compare the behavior of 3xHA-tagged SNC1 introduced into *snc1-11* and *adr1-L1-c1/snc1-11* plants. Upon inactivation of *ADR1-L1*, SNC1 dramatically shifts to a slow-migrating species of 480-720 kDa, which likely corresponds to SNC1 tetramers **(Fig. 6a)**. We conclude that the loss of ADR1-L1 is sufficient to trigger the oligomerization of SNC1, with the SNC1 oligomer constituting the active form.

**Fig. 6.**
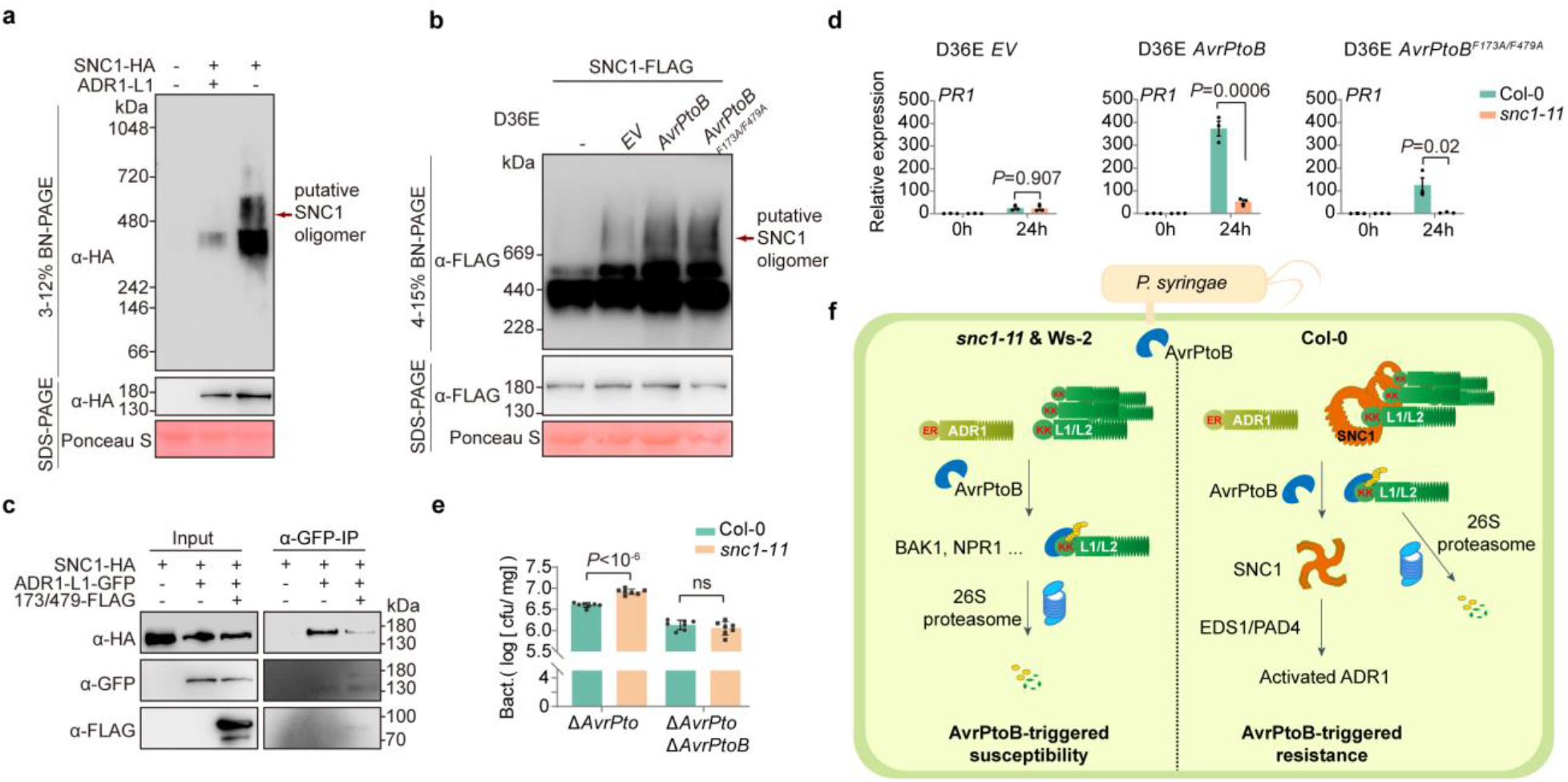
AvrPtoB induces oligomerization of SNC1 and activates SNC1-dependent immune responses. **a**, Absence of ADR1-L1 stimulates SNC1 oligomerization in Arabidopsis, as shown by BN-PAGE and SDS-PAGE. Arrow points to apparent higher-order SNC1 complexes, likely SNC1 tetramers. **b**, AvrPtoB enhances SNC1 oligomerization as shown by BN-PAGE and SDS-PAGE. Arrow points to potential SNC1 complexes. **c**, The E3 ligase dead variant AvrPtoB^F173A/F479A^ interferes with the interaction between ADR1-L1 and SNC1, as shown by Co-IP assays in *N. benthamiana*. **d**, *PR1* expression of Col-0 and *snc1-11* infiltrated with *Pst* DC3000 D36E carrying empty vector, AvrPtoB and AvrPtoB^F173A/F479A^, as measured by RT-qPCR. Data represent the mean and standard error of three biological replicates (n = 3 biologically independent samples, *p*-values from Student’s t-test). **e**, AvrPtoB activates SNC1-dependent resistance to *Pst*. Bacterial growth assays of *Pst* DC3000 *ΔAvrPto* and *Pst* DC3000 *ΔAvrPto ΔAvrPtoB* on Col-0 and *snc1-11* (n = 3 biologically independent samples, *p*-values from Student’s t-test). **(a-e)** Experiments were performed three times, with similar results. **f**, Working model. In absence of functional SNC1, for example, in Ws-2 and *snc1-11*, AvrPtoB ubiquitinates ADR1-L1, and to a lesser extent ADR1-L2, to promote their degradation, preventing activation of immunity. In the presence of SNC1, degradation of ADR1-L1 and ADR1-L2 induced by AvrPtoB activates oligomerization of the guarding NLR SNC1, which relays signals through ADR1 to trigger downstream immune responses.

Since ubiquitination of ADR1-L1 by AvrPtoB leads to its removal, akin to the situation in *adr1-L1-c1* mutants, we also examined whether AvrPtoB induced SNC1 oligomerization. As shown in **Fig. 6b**, infiltration of *Pst* DC3000 D36E expressing AvrPtoB induced a slow-migrating SNC1 species of 480-720 kDa, similar to what had been observed in *adr1-L1-c1* mutants **(Fig. 6a)**, confirming that SNC1 acts as a guard for the AvrPtoB target ADR1-L1. Unexpectedly, infiltration of *Pst* DC3000 D36E carrying the E3 ligase dead AvrPtoB^F173A/F479A^ also triggered a slow-migrating SNC1 species of 480-720 kDa. Since both SNC1 and AvrPtoB interact with the CC_R_ domain of ADR1-L1, AvrPtoB^F173A/F479A^ may compete with the binding of ADR1-L1 to SNC1, which would result in failure of ADR1-L1 to prevent oligomerization of SNC1. To test this hypothesis, ADR1-L1-GFP and SNC1-HA were co-expressed with AvrPtoB^F173A/F479A^-FLAG for Co-IP assays in *N. benthamiana*. In support of the proposed scenario, AvrPtoB^F173A/F479A^ substantially reduced the ability of ADR1-L1 to pull down SNC1 **(Fig. 6c)**.

Overexpression of AvrPtoB induces dramatic autoimmunity in the Col-0 accession^36^, which we hypothesized could be due to loss of ADR1-L1 and concomitant activation of SNC1. Attempts to generate *35S::AvrPtoB-FLAG* transgenic lines for epistasis analysis with *SNC1* were not successful, likely due to extreme autoimmunity. As alternative, we measured expression of the defense marker *PR1* in Arabidopsis plants upon delivery of AvrPtoB or AvrPtoB^F173A/F479A^ by DC3000 D36E.

As shown in **Fig. 6d**, *snc1-11* mutants expressed significantly less *PR1* than wild-type plants in these trials. Moreover, the higher growth of *Pst* DC3000 *ΔAvrPto* in *snc1-11* mutants compared to wild-type plants was dependent on AvrPtoB since no difference was seen between *snc1-11* and wild-type plants infiltrated with *Pst* DC3000 *ΔAvrPto ΔAvrPtoB* **(Fig. 6e)**. We conclude that the degradation of ADR1-L1 initiated by AvrPtoB activates immune responses mediated by SNC1.

## DISCUSSION

The conserved helper NLR proteins of the ADR1 family are key ETI components^17^. We found that the bacterial effector AvrPtoB targets ADR1 homologs, and that these are in turn guarded by the sensor NLR SNC1. Our findings demonstrate a new concept in the tug of war between pathogens using effectors and plants using immune receptors, and they reveal also the long-sought after function of SNC1 in plant immunity.

Because NLR over-accumulation can trigger spontaneous autoimmunity, NLR abundance is tightly controlled at multiple levels^17^. For example, to maintain NLR protein homeostasis, plants evolved a set of E3 ubiquitin ligases to regulate NLR stability. The plant E3 ligases CPR1/CPR30 and SNIPER1/2 ubiquitinate SNC1, thereby limiting SNC1 levels, and their knockout triggers SNC1-mediated autoimmunity^37^. Here, we show that *Pseudomonas* utilizes in a similar manner the E3 ligase AvrPtoB effector to induce degradation of the helper NLRs ADR1-L1/2, but in this case reduced NLR protein levels lead to autoimmunity because ADR1-L1/2 is a client for the sensor NLR SNC1.

AvrPtoB is a conserved effector found in the genomes of diverse Gram-negative bacteria, including *Pseudomonas, Xanthomonas* and *Erwinia*^38^. AvrPtoB has been shown to target and ubiquitinate a wide range of proteins, including several pattern recognition receptors and PTI key component BAK1 (BRASSINOSTEROID RECEPTOR-ASSOCIATED KINASE 1)^39^, the master regulator of salicylic acid signalling, NPR1 (NON-EXPRESSER OF PR GENES 1)^40^, and an exocyst subunit^36^. Here we show that AvrPtoB can dampen both PTI and ETI, by identifying the central ETI components ADR1-L1 and ADR-L2 as AvrPtoB targets.

Pathogen effectors have two roles: One is to manipulate host physiology for the colonizer’s benefit, the other – and the one most recent work has focused on – is to suppress host defences, especially those related to PTI^41^. PTI and ETI are inter-linked ^3–6^, and the targeting of PTI versus ETI by effectors cannot always be neatly separated. EDS1 was initially identified as a key ETI component, forming EDS1-PAD4-ADR1 and EDS1-SAG101-NRG complexes that regulate transcriptional reprogramming during defence and HR^42^. The EDS1-PAD4-ADR1 module plays, however, also an important role in PTI^3,6^.

Examples of effector targeting NLR come from the *P. infestans* effector AVRcap1b and the cyst nematode effector SS15, which suppress Solanaceae-specific helper NLRs NRC2 and NRC3 either by affecting their negative regulator NbTOL9a or by preventing their oligomerization and activation^22,23^. We add to these insights, by revealing not only that helper NLRs ADR1-L1 and ADR1-L2 are targeted by *P. syringae* effector AvrPtoB, but also that AvrPtoB-induced degradation of ADR1-L1 and ADR1-L2 is monitored by the sensor NLR SNC1 **(Fig. 6f)**. Effectors of independent origin often converge on conserved targets with essential roles in plant immunity^43^. ADR1 homologs, which are widespread in the plant kingdom^7^, clearly fulfil this definition, and it is therefore not unlikely that other effectors targeting ADR1 homologs await discovery. Similarly, it will be of interest to learn whether ADR1 homologs in other species are guarded by NLRs as well, and whether such interactions mimic the interaction between ADR1-L1/L2 and SNC1 in Arabidopsis.

One of the reasons that there is a rich literature on SNC1 is that its knockout suppresses, albeit to different degrees, autoimmunity resulting from changes in a wide range of proteins^17^. Given the role of SNC1 as a guard of ADR1 homologs, the genetic interactors of SNC1 might be negative regulators of SNC1, potentially by affecting the interaction between SNC1 and ADR1 homologs. Guarding of ADR1 homologs might, however, not be the only role of SNC1, which has been proposed to be a more general amplifier of ETI^44^. SNC1 was found to enhance avrRpt2- and avrRps4-induced resistance^44^, which depends on ADR1 homologs^45^. We propose that the formation of ADR1 oligomers triggered by interaction of effectors such as AvrRpt2 and AvrRps4 with their cognate NLR immune receptors could displace SNC1 from the ADR-L1/2-SNC1 guardee-guard complex, which in turn might amplify downstream immune responses via ADR1. Regardless of any other roles, however, SNC1 clearly fits the definition of a resistance protein for indirect recognition of the bacterial effector AvrPtoB. The importance of being able to detect AvrPtoB is also apparent from the fact that, as with other effectors^17^, AvrPtoB can be recognized by other NLRs, including tomato Prf via its guardee Pto, which directly interacts with AvrPtoB^1,47^.

In summary, we have demonstrated that bacterial AvrPtoB ubiquitinates conserved key components of ETI, which in turn is detected by the plant host through the sensor NLR SNC1. Our work highlights how the same pathway can be a target of pathogen effector proteins and at the same time be used to protect the host from these effectors. In addition, we demonstrate how sequence diversification enables a partially redundant helper NLR to evade effector suppression and thereby preserve the integrity of ETI.

## METHODS

### Plant material and growth conditions

*Arabidopsis thaliana* and *Nicotiana benthamiana* were derived from stocks maintained in the lab. Arabidopsis mutants and transgenic plants generated in this study are listed in the key resource table. Arabidopsis plants were grown under long-day (16 h day/8 h night) or short-day (10 h day/14 h night) regimes at 23°C with relative humidity at 65%. *Nicotiana benthamiana* plants were grown in a greenhouse under long-day conditions for 4-5 weeks before transient transformation.

### Cell death assays

For the cell death assays, autoactive variants of ADR1s were co-expressed with indicated genes in *N. benthamiana* through agroinfiltration. Briefly, *Agrobacterium tumefaciens* GV3101 containing the relevant expression vectors were grown in liquid LB(Lysogeny broth) medium overnight in a shaking incubator (220 rpm, 28°C). *Agrobacteria* were precipitated through centrifugation and re-suspended in an infiltration buffer (10 mM MgCl_2_, 10 mM MES, pH 5.6). Vectors used for cell death assays are listed in **Supplementary Table 1**. For co-expression, each bacterial suspension was adjusted to the final OD_600_ indicated in **Supplementary Table 1**, and infiltrated into 4-week-old *N. benthamiana* plants. The HR phenotypes were photographed and scored 2-3 days after agroinfiltration.

### Generation of transgene-free gene-edited lines

The gRNA sequences, gRNA1 5’-GAGCTCCATTGACTTGACT-3’, gRNA2 5’-CTATAACGTTAACCGGTAG-3’, and gRNA3 5’-GCTCACCGCCAAGCTCAAAT-3’ were introduced to pREE401E, which was modified from an egg cell-specific CRISPR-CAS9 toolkit vector pHEE401E by adding Fast-RED selection marker^48,49^, to knock-out *ADR1-L1*. The gene editing events were verified by PCR and Sanger sequencing. T_2_ seeds that without red fluorescent seed coats were isolated as transgene-free seeds.

### Generation of high-order mutants

To generate high-order mutants, *adr1-L1-c1* was crossed with *pad4-1, sid2-2, ndr1-1, eds1-2, nrg* triple, *adr1* triple. The homozygous high-order mutants were verified by PCR or Sanger sequencing. The genotyping primers are listed in **Supplementary Table 2**.

### RT-qPCR

RNA was extracted from plant tissue using an RNA isolation method (R401, Vazyme Biotech Co. Ltd. Nanjing, China). cDNA was synthesized from 0.5 μg high-quality total RNA (A260/A230>2.0 and A260/A280>1.8), using HiScript Ⅲ First Strand cDNA Synthesis (R312, Vazyme Biotech Co. Ltd. Nanjing, China). SYBR master mix (Q711, Vazyme Biotech Co. Ltd., Nanjing, China) was used for quantitative real-time PCR in a Thermo Fisher system (ABI QuantStudio 6 Flex) according to the manufacturer’s instructions. The comparative Ct (ΔΔCt) method was used to calculate the relative expression of genes of interest, using *ACTIN2* gene (*AT3G18780*) as an internal control. The primers used for qPCR are listed in **Supplementary Table 2**.

### Phylogeny analysis

To construct the phylogenetic tree of ADR1 homologs in angiosperms, the amino acid sequence of CC_R_ ^ADR1-L1^ was used as query to BLAST in NCBI. The resulted sequences, which feature typical CC_R_, NB-ARC, and LRR domains, were used for further analysis. The MAFFT aligned sequences of the NB-ARC domain were used for phylogeny analysis with PhyML in NGPhylogeny.fr webserver^50^. Sequence LOGOs of ADR1, ADR1-L1, and ADR1-L2 in Brassicaceae were created by WebLOGO webserver^51^ with grouped sequences according to phylogeny analysis results.

### Constructs and transgenic lines

The genomic fragments of *ADR1, ADR1-L1*, and *ADR1-L2* were amplified through PCR using Col-0 genomic DNA as template. The resulting PCR products were cloned into entry vector pUC19 using homologous recombination (C115, Vazyme Biotech Co. Ltd. Nanjing, China) and transferred into the binary vector pCambia1300, which contains hygromycin marker for plant selection. To generate *pT-DNA::ADR1-L1*^*Δ12aa*^ and *p35S::ADR1-L1*^*Δ12aa*^, the truncated *ADR1-L1*^*Δ12aa*^ CDS fragment was amplified from cDNA of SAIL_302_C06, and a 2 kb of T-DNA fragment near to insertion site and *35S CaMV* fragment were amplified as promoters for *ADR1-L1*^*Δ12aa*^. The corresponding promoter and the *ADR1-L1*^*Δ12aa*^ amplicon were cloned into pCambia1300 by multiple fragments homologous recombination. The CDS of *CC*^*RADR1s*^ were amplified from Col-0 cDNA, cloned into the entry vector pUC19, and then subcloned into the binary vector pCBNS-GFP. The CDS of AvrPtoB was amplified using *Pst* DC3000 genomic DNA and cloned into pCBCS-HA/-FLAG and pME6012 by homologous recombination.

Site-directed mutagenesis and chimeric constructs were carried out by introducing corresponding changes in the primers using multiple fragments homologous recombination.

Primer sequences used for domain swap and site-directed mutagenesis were listed in **Supplementary Table 2**.

The expression constructs were introduced into *Agrobacterium tumefaciens* GV3101 by electroporation. Stable transgenic plants were generated through the floral dipping method^52^. T_1_ transformants were screened based on hygromycin selection or red fluorescent selection.

### Map-based cloning

To map the natural suppressor(s) of *adr1-L1* in Ws-2, a F_2_ mapping population derived from a cross between *adr1-L1*^*Ws-2*^ and *adr1-L1-c1* was generated. F_2_ individuals with normal growth phenotypes were selected for genotyping. The SSLP markers were designed according to Yang’s previous work^31^, and the detailed information is provided in **Supplementary Table 2**.

### Bacterial infection

For the bacterial infection assays on soil-grown plants, *Pst* DC3000 was precipitated by centrifugation and suspended in 10mM MgCl_2_ solution. The concentrations of *Pst* DC3000 were adjusted to OD600 = 0.002. *Pst* DC3000 was infiltrated into rosette leaves with a needleless syringe. Leaf discs (6 mm) from inoculated leaves were collected at 3 dpi.

For the bacterial infection assays on germ-free plants, seedlings were grown on 1/2 Murashige and Skoog (MS) medium in 90 × 90 mm culture plate for three weeks. Bacteria were grown overnight at 28°C in the King’s B medium plates with appropriate antibiotics. Bacteria were harvested from the plates, resuspended in sterile water with 0.025% Silwet L-77, and the concentration of *Pst* DC3000 Δ*AvrPto* and *Pst* DC3000 Δ*AvrPt*o Δ*AvrPtoB* were adjusted to an optical density at OD600 = 0.02. 50 ml of bacterial suspension was poured onto the culture plates containing 3-week-old plant and rested for 3 min at room temperature. After removing the bacterial suspension by decantation, the plates were sealed with 3M Micropore surgical tape and incubated at the growth chamber. The whole plant was weighed and collected at 2 dpi.

### AvrPtoB-induced protein degradation in Arabidopsis

For the protein degradation assays, Arabidopsis plants were grown under short-day conditions. *Pseudomonas syringae* DC3000 D36E strains containing *EV, AvrPto*B, or *AvrPtoB*^*F173A/F479A*^, were cultured on solid KB (King’s B) medium at 28°C for 24 hours. Bacterial suspensions were adjusted to an OD600 of 0.4 in 10 mM MgCl2 solution, then infiltrated into 4-week-old Arabidopsis plants with a needleless syringe. Leaf discs at a diameter of 6 mm were collected from inoculated leaves at 0 hpi, 3 hpi, 6 hpi, and 12 hpi for immunoblots.

### Split-luciferase complementation assay

In the Split-Luc assays, AvrPtoB-nLuc was transiently co-expressed with ADR1-cLuc, ADR1-L1-cLuc, ADR1-L2-cLuc, and EV in 4-week-old *N. benthamiana* leaves. At 2 days post-infiltration (dpi) with *Agrobacterium* strains harbouring the relevant constructs, leaves were infiltrated with 1 mM luciferin containing 0.02% Silwet L-77 and kept in the dark for 5 minutes before CCD imaging. To quantify the luciferase signal, leaf discs were collected from the inoculated leaves using a 6 mm puncher and placed into a 96-well plate with 60 μl H_2_O. 60 μl of 2 mM luciferin was added to the leaf discs in the 96-well plate before recording luminescence.

### Co-immunoprecipitation

*Agrobacterium* strains harbouring AvrPtoB-HA, SNC1-H, ADR1-FLAG, ADR1-L1-FLAG, and ADR1-L2-FLAG were grown overnight in LB medium containing appropriate antibiotics (220 rpm, 28°C) and used for agroinfiltration in *N. benthamiana*. Inoculated leaves were harvested 2dpi and ground into powder with liquid nitrogen. Ground tissues were homogenized in ice-cold extraction buffer (10% glycerol, 25 mM Tris-HCl pH 7.5, 1 mM EDTA, 150 mM NaCl, 2% PVP, 0.5% Triton-X100) supplemented with 1 mM DTT, anti-protease tablet (04693132001, Roche, USA). The resulting lysate was homogenized by mixing for 20 min on ice and centrifuged at 13000 rpm for 15 min at 4°C, with this step being repeated twice. The supernatant was incubated with 5 μl Antibodies-coupled beads (Anti-FLAG M2, M8823, Sigma-Aldrich, USA; Anti-GFP, KTSM1334, KangTi Life Technology, Shenzhen, China) for 3 hours at 4°C under gentle agitation. After incubation, beads were washed six times with washing buffer (25 mM Tris-HCl pH 7.5, 1 mM EDTA, 150 mM NaCl, 0.5% Triton-X 100,1 mM DTT) at 4°C.

SDS-loading buffer (8 M urea, 2% SDS, 20% glycerol, 100 mM Tris-HCl pH 6.8, 0.004% bromophenol blue) with 100 mM DTT was added to beads before boiling at 95°C for 5 min to release bound proteins. Released proteins were analysed by immunoblots.

### In vitro ubiquitination assays

Bacteria (BL21) harbouring GST-, MBP-6xHis-, and 6xHis-fusion protein expression vectors were cultured in LB at 37°C until an OD_600_ of 0.6. Protein expression was induced by adding 0.4 mM IPTG and incubating at 16°C for 16 hours. Tagged proteins were purified separately using Glutathione Sepharose 4B (17075601, GE Healthcare, Chicago, USA) or Ni-NTA affinity agarose beads (30210, QIAGEN, Venlo, Netherlands).

Ubiquitination reactions were performed in a total volume of 30 μl, consisting of 50 mM Tris-HCl (pH 7.5), 2 mM ATP, 1 mM MgCl_2_, 1 mM DTT, 500 mg E1-His, 1 μg E2-His, 3 μg GST-AvrPtoB, 500ng MBP-CC_R_s and 3 μg ubiquitin for 8 h at 30 °C. Reactions were stopped by adding 30 μl SDS-loading buffer (8 M urea, 2% SDS, 20% glycerol, 100 mM Tris-HCl pH 6.8, 0.004% bromophenol blue) and the samples were boiled for 5 min at 95°C.

### In vitro pull-down assays

For the GST pull-down assays, 2 µg GST-tagged Protein, 20 µl Glutathione Sepharose 4B (17075601, GE Healthcare, Chicago, USA) and 10 µg MBP-6xHis-tagged protein were added to 1 ml pull-down buffer (50 mM Tris-HCl [pH 7.5], 200 mM NaCl, 0.5% [v/v] Triton X-100) and incubated for 4 hours under gentle rotation. Beads were washed 6 times with 1 ml pull-down buffer. SDS-loading buffers were added to beads before boiling to release bound proteins. The released proteins were analysed by immunoblots using anti-Glutathione-S-Transferase (AE001, AbClonal, Wuhan, China) and anti-MBP (AE016, AbClonal, Wuhan, China) antibodies.

For the SNC1-GFP pull-down assays, ground *N. benthamiana* leaves transiently expressing SNC1-GFP were homogenized in extraction buffer containing 10% glycerol, 25 mM Tris-HCl pH 7.5, 1 mM EDTA, 150 mM NaCl, 2% PVP, 0.5% Triton-X 100, 1 mM DTT, and protease inhibitor. The resulting lysate was centrifuged and subjected for SNC1-GFP precipitation using anti-GFP magnetic beads (KTSM1334, KangTi Life Technology, Shenzhen, China). The anti-GFP magnetic beads were then aliquoted into 4 tubes containing 2 μg MBP-tagged protein in 1 ml buffer containing 25 mM Tris-HCl pH 7.5, 1 mM EDTA, 150 mM NaCl, and 0.5% Triton-X100, and incubated for 3 hours under gentle rotation. Beads are washed 6 times with 1 ml pull-down buffer. SDS-loading buffers were added to beads before boiling to release bound proteins. The released proteins were analysed by immunoblots using anti-GFP (AE012, AbClonal, Wuhan, China) and anti-MBP (AE016, AbClonal, Wuhan, China) antibodies.

### Blue Native-PAGE

Blue native polyacrylamide gel electrophoresis (BN-PAGE) was performed according to ref. 53. Three 14-day-old seedlings, infected with or without *Pst* D36E, were collected and homogenized in 1 x NativePAGE Sample Buffer (BN20032, Invitrogen, CA, USA) supplemented with 1% n-dodecyl β-D-maltoside (DDM) and protease inhibitor cocktail (4693116001, Roche, USA). Homogenization was achieved by gently mixing on ice for 20 min, followed by 20000 g centrifugation for 15 min at 4℃. The resulting supernatant was mixed with 0.25% G-250 Sample Additive and loaded on a NativePAGE 3-12% Bis-Tris gel (BN1001BOX, Invitrogen, CA, USA) for electrophoresis.

### Data availability

This study analyses existing, publicly available sequencing data and does not disclose new datasets and sequences. All data are provided in the main figures and extended data.

## ACKNOWLEDGEMENTS

We thank Lei Li (CAS), Guozhi Bi (CAU), Yule Liu (THU) for discussion. We thank Wenbo Ma (TSL) for critical reading of the manuscript. We thank Lei Li and He Zhao (TSL) for technical support with the BN-PAGE experiment. We thank Xin Li (UBC) for helper-NLR mutant seeds, Hailei Wei (CAAS) for the *Pst* DC3000 D36E strain, Jun Liu (CAU) for the *Pst* DC3000 T3SSS effector vector library and the DC3000 *ΔAvrPto* strain, and Fuhao Cui (CAU) for the *Pst* DC3000 *ΔAvrPto ΔAvrPtoB* strain. R.W. was supported by the EU Horizon 2020 research and innovation programme under the Marie Skłodowska-Curie scheme (H2020-MSCA-IF-2014-655295). J.D.J. was supported by the Gatsby Foundation (UK). D.W. was supported by the Max Planck Society.

J.H. was supported by the EUs Horizon 2020 research and innovation programme under the Marie Skłodowska-Curie scheme (No 897584). W.Z. was supported by the National Key Research and Development Program, Ministry of Science and Technology of China (No 2022YFD1201802), the Ministry of Education of China (the 111 Project B13006) and the 2115 Talent Development Program of China Agricultural University (No 2020RC013).

## AUTHOR CONTRIBUTIONS

Conceptualization: J.H., W.Z. Methodology: M.W., J.C., R.W., H.G., H.G., J.H., W.Z. Formal analysis: M.W., J.C., R.W., W.Z. Investigation: M.W., J.C., R.W., H.G., Y.C., Z.L., L.W., C.L., S.H., M.D., H.G. Writing–original draft: M.W., J.C., J.H., W.Z. Writing–review, and editing: J.C., J.H., W.Z., D.W. Supervision: Y.P., D.W., J.D.J., W.Z. Project administration: W.Z. Funding acquisition: R.W., J.D.J., D.W., J.H., W.Z.

## REFERENCES

1. Jones, J. D. G. & Dangl, J. L. The plant immune system. Nature 444, 323–329 (2006).

2. Zhou, J.-M. & Zhang, Y. Plant Immunity: Danger Perception and Signaling. Cell 181, 978–989 (2020).

3. Pruitt, R. N. et al. The EDS1–PAD4–ADR1 node mediates Arabidopsis pattern-triggered immunity. Nature 1–5 (2021).

4. Yuan, M. et al. Pattern-recognition receptors are required for NLR-mediated plant immunity. Nature 592, 105–109 (2021).

5. Ngou, B. P. M., Ahn, H.-K., Ding, P. & Jones, J. D. G. Mutual potentiation of plant immunity by cell-surface and intracellular receptors. Nature 592, 110–115 (2021).

6. Tian, H. et al. Activation of TIR signalling boosts pattern-triggered immunity. Nature 598, 500–503 (2021).

7. Liu, Y. et al. An angiosperm NLR atlas reveals that NLR gene reduction is associated with ecological specialization and signal transduction component deletion. Mol. Plant 14, 2015–2031 (2021).

8. Bi, G. et al. The ZAR1 resistosome is a calcium-permeable channel triggering plant immune signaling. Cell 184, 3528–3541.e12 (2021).

9. Förderer, A. et al. A wheat resistosome defines common principles of immune receptor channels. Nature 610, 532–539 (2022).

10. Jacob, P. et al. Plant ‘helper’ immune receptors are Ca2+-permeable nonselective cation channels. Science 373, 420–425 (2021).

11. Huang, S. et al. Identification and receptor mechanism of TIR-catalyzed small molecules in plant immunity. Science 377, eabq3297 (2022).

12. Wu, Z. et al. Differential regulation of TNL-mediated immune signaling by redundant helper CNLs. New Phytol. 222, 938–953 (2019).

13. Castel, B. et al. Diverse NLR immune receptors activate defence via the RPW8-NLR NRG1. New Phytol. 222, 966–980 (2019).

14. Zhang, Y., Goritschnig, S., Dong, X. & Li, X. A gain-of-function mutation in a plant disease resistance gene leads to constitutive activation of downstream signal transduction pathways in suppressor of npr1-1, constitutive 1. Plant Cell 15, 2636–2646 (2003).

15. Stokes, T. L., Kunkel, B. N. & Richards, E. J. Epigenetic variation in Arabidopsis disease resistance. Genes Dev. 16, 171–182 (2002).

16. Li, X., Clarke, J. D., Zhang, Y. & Dong, X. Activation of an EDS1-mediated R-gene pathway in the snc1 mutant leads to constitutive, NPR1-independent pathogen resistance. Mol. Plant. Microbe. Interact. 14, 1131–1139 (2001).

17. van Wersch, S., Tian, L., Hoy, R. & Li, X. Plant NLRs: The Whistleblowers of Plant Immunity. Plant Commun 1, 100016 (2020).

18. Zhu, W. et al. Modulation of ACD6 dependent hyperimmunity by natural alleles of an Arabidopsis thaliana NLR resistance gene. PLoS Genet. 14, e1007628 (2018).

19. Wei, H.-L. et al. Pseudomonas syringae pv. tomato DC3000 Type III Secretion Effector Polymutants Reveal an Interplay between HopAD1 and AvrPtoB. Cell Host Microbe 17, 752–762 (2015).

20. Martel, A. et al. Metaeffector interactions modulate the type III effector-triggered immunity load of Pseudomonas syringae. PLoS Pathog. 18, e1010541 (2022).

21. Wei, H.-L., Zhang, W. & Collmer, A. Modular Study of the Type III Effector Repertoire in Pseudomonas syringae pv. tomato DC3000 Reveals a Matrix of Effector Interplay in Pathogenesis. Cell Rep. 23, 1630–1638 (2018).

22. Derevnina, L. et al. Plant pathogens convergently evolved to counteract redundant nodes of an NLR immune receptor network. PLoS Biol. 19, e3001136 (2021).

23. Contreras, M. P. et al. Resurrection of plant disease resistance proteins via helper NLR bioengineering. bioRxiv 2022.12.11.519957 (2022) doi:10.1101/2022.12.11.519957.

24. Janjusevic, R., Abramovitch, R. B., Martin, G. B. & Stebbins, C. E. A bacterial inhibitor of host programmed cell death defenses is an E3 ubiquitin ligase. Science 311, 222–226 (2006).

25. Collier, S. M., Hamel, L.-P. & Moffett, P. Cell death mediated by the N-terminal domains of a unique and highly conserved class of NB-LRR protein. Mol. Plant. Microbe. Interact. 24, 918–931 (2011).

26. Bonardi, V. et al. Expanded functions for a family of plant intracellular immune receptors beyond specific recognition of pathogen effectors. Proc. Natl. Acad. Sci. U. S. A. 108, 16463–16468 (2011).

27. Dong, O. X. et al. TNL-mediated immunity in Arabidopsis requires complex regulation of the redundant ADR1 gene family. New Phytol. 210, 960–973 (2016).

28. van Wersch, R., Li, X. & Zhang, Y. Mighty Dwarfs: Arabidopsis Autoimmune Mutants and Their Usages in Genetic Dissection of Plant Immunity. Front. Plant Sci. 7, 1717 (2016).

29. Cui, H. et al. A core function of EDS1 with PAD4 is to protect the salicylic acid defense sector in Arabidopsis immunity. New Phytol. 213, 1802–1817 (2017).

30. Van de Weyer, A.-L. et al. A Species-Wide Inventory of NLR Genes and Alleles in Arabidopsis thaliana. Cell 178, 1260–1272.e14 (2019).

31. Yang, S. & Hua, J. A haplotype-specific Resistance gene regulated by BONZAI1 mediates temperature-dependent growth control in Arabidopsis. Plant Cell 16, 1060–1071 (2004).

32. Wang, J. et al. Reconstitution and structure of a plant NLR resistosome conferring immunity. Science 364, eaav5870 (2019).

33. Wang, J. et al. Ligand-triggered allosteric ADP release primes a plant NLR complex. Science 364, eaav5868 (2019).

34. Martin, R. et al. Structure of the activated ROQ1 resistosome directly recognizing the pathogen effector XopQ. Science 370, (2020).

35. Ma, S. et al. Direct pathogen-induced assembly of an NLR immune receptor complex to form a holoenzyme. Science 370, (2020).

36. Wang, W., Liu, N., Gao, C., Rui, L. & Tang, D. The Pseudomonas Syringae Effector AvrPtoB Associates With and Ubiquitinates Arabidopsis Exocyst Subunit EXO70B1. Front. Plant Sci. 10, 1027 (2019).

37. Lapin, D., Johanndrees, O., Wu, Z., Li, X. & Parker, J. E. Molecular innovations in plant TIRbased immunity signaling. Plant Cell 34: 1479–1496 (2022).

38. Abramovitch, R. B., Kim, Y.-J., Chen, S., Dickman, M. B. & Martin, G. B. Pseudomonas type III effector AvrPtoB induces plant disease susceptibility by inhibition of host programmed cell death. EMBO J. 22, 60–69 (2003).

39. Wei, H.-L. & Collmer, A. Defining essential processes in plant pathogenesis with Pseudomonas syringae pv. tomato DC3000 disarmed polymutants and a subset of key type III effectors. Mol. Plant Pathol. 19, 1779–1794 (2018).

40. Chen, H. et al. A Bacterial Type III Effector Targets the Master Regulator of Salicylic Acid Signaling, NPR1, to Subvert Plant Immunity. Cell Host Microbe 22, 777–788.e7 (2017).

41. Deslandes, L. & Rivas, S. Catch me if you can: bacterial effectors and plant targets. Trends Plant Sci. 17, 644–655 (2012).

42. Sun, X. et al. Pathogen effector recognition-dependent association of NRG1 with EDS1 and SAG101 in TNL receptor immunity. Nat. Commun. 12, 3335 (2021).

43. Weßling, R. et al. Convergent targeting of a common host protein-network by pathogen effectors from three kingdoms of life. Cell Host Microbe 16, 364–375 (2014).

44. Wang, Z., Yang, L. & Hua, J. The intracellular immune receptor like gene SNC1 is an enhancer of effector-triggered immunity in Arabidopsis. Plant Physiol. 191, 874–884 (2022).

45. Saile, S. C. et al. Two unequally redundant ‘helper’ immune receptor families mediate Arabidopsis thaliana intracellular ‘sensor’ immune receptor functions. PLoS Biol. 18, e3000783 (2020).

46. Mucyn, T. S. et al. The tomato NBARC-LRR protein Prf interacts with Pto kinase in vivo to regulate specific plant immunity. Plant Cell 18, 2792–2806 (2006).

47. Mathieu, J., Schwizer, S. & Martin, G. B. Pto kinase binds two domains of AvrPtoB and its proximity to the effector E3 ligase determines if it evades degradation and activates plant immunity. PLoS Pathog. 10, e1004227 (2014).

## REFERENCES

48. Shimada, T. L., Shimada, T. & Hara-Nishimura, I. A rapid and non-destructive screenable marker, FAST, for identifying transformed seeds of Arabidopsis thaliana. Plant J. 61, 519–528 (2010).

49. Wang, Z.-P. et al. Egg cell-specific promoter-controlled CRISPR/Cas9 efficiently generates homozygous mutants for multiple target genes in Arabidopsis in a single generation. Genome Biol. 16, 144 (2015).

50. Lemoine, F. et al. NGPhylogeny.fr: new generation phylogenetic services for non-specialists. Nucleic Acids Res. 47, W260–W265 (2019).

51. Crooks, G. E., Hon, G., Chandonia, J.-M. & Brenner, S. E. WebLogo: a sequence logo generator. Genome Res. 14, 1188–1190 (2004).

52. Clough, S. J. & Bent, A. F. Floral dip: a simplified method for Agrobacterium-mediated transformation of Arabidopsis thaliana. Plant J. 16, 735–743 (1998).

53. Na Ayutthaya, P. P., Lundberg, D., Weigel, D. & Li, L. Blue Native Polyacrylamide Gel Electrophoresis (BN-PAGE) for the Analysis of Protein Oligomers in Plants. Curr. Protoc. Plant Biol. 5, e20107 (2020).

