## Supplemental Figures for "The immune receptor SNC1 monitors helper NLRs targeted by a bacterial effector"

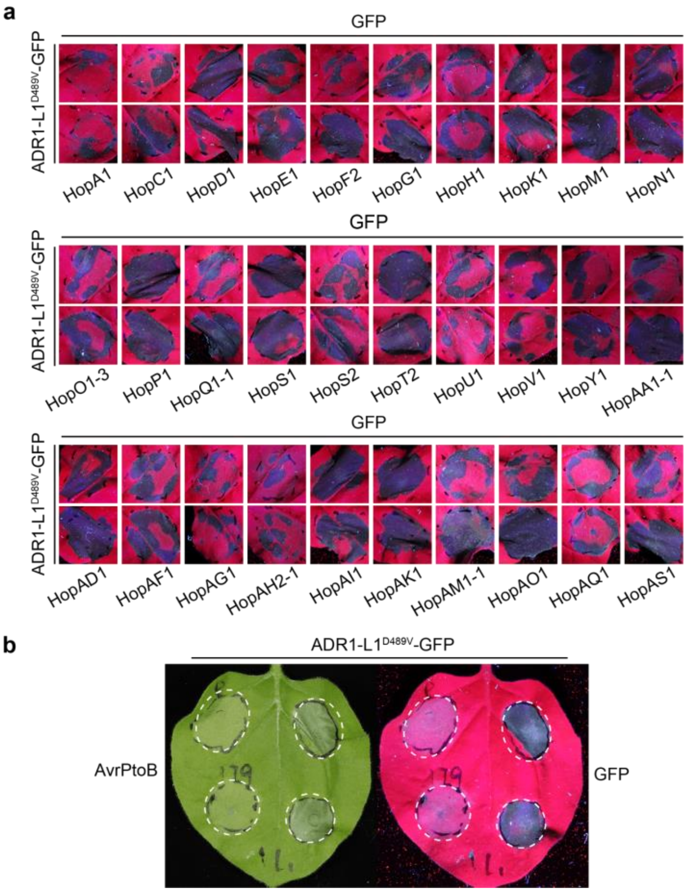


**Extended Data Fig. 1. Identification of *Pst* DC3000 T3SS effectors that suppress ADR1-L1-triggered HR in *N. benthamiana*. a,** A construct for autoactivate ADR1-L1$^{D489V}$ was transformed into *Agrobacterium tumefaciens* and transiently co-expressed in *N. benthamiana* plants with each effector or GPF-HA control vector. Thirty-one T3SS effectors were tested. **b,** Representative *N. benthamiana* leaves showing ADR1-L1$^{D489V}$-triggered HR suppressed by co-expression of AvrPtoB.

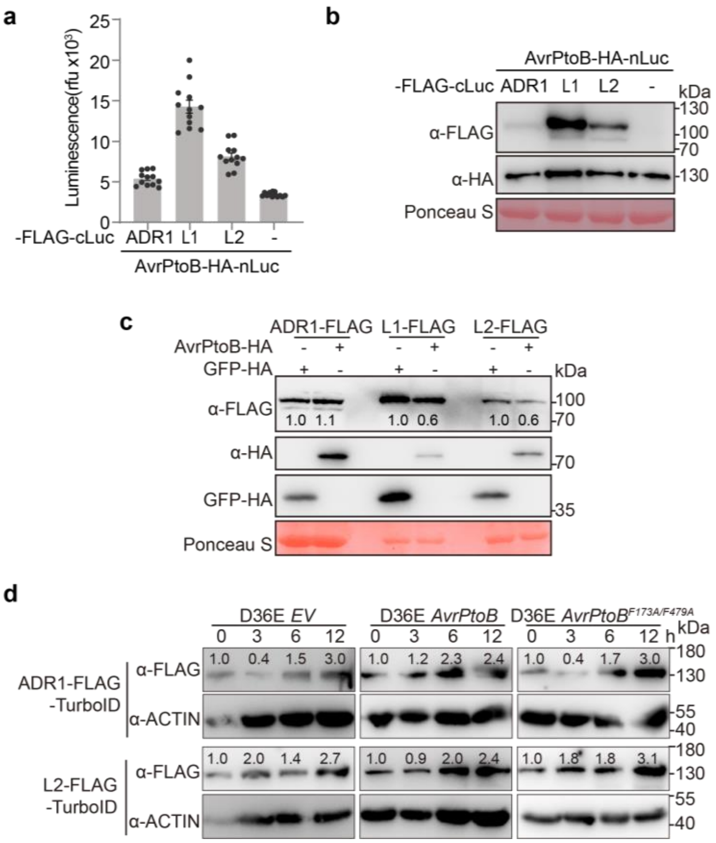


**Extended Data Fig. 2. Requirement of ADR1-L1 CC$_R$ domain for AvrPtoB-mediated suppression of ADR1-L1$^{D489V}$-triggered HR. a,** Quantitative measurement of luciferase signals showing that AvrPtoB interacts with ADR1-L1 more strongly than with ADR1 and ADR1-L2. **b,** Protein expression in split-luciferase complementation assays shown in **(a)**. **c,** AvrPtoB reduces accumulation of Flag-tagged ADR1-L1 and ADR1-L2 but not ADR1 protein in *N. benthamiana*. **d,** AvrPtoB does not reduce accumulation of ADR1-FLAG-TurboID and ADR1-L2-FLAG-TurboID in transgenic Arabidopsis plants. Numbers on top of the gel blots indicate arbitrary densitometry units of corresponding bands after normalization to the sample collected at 0 hour.

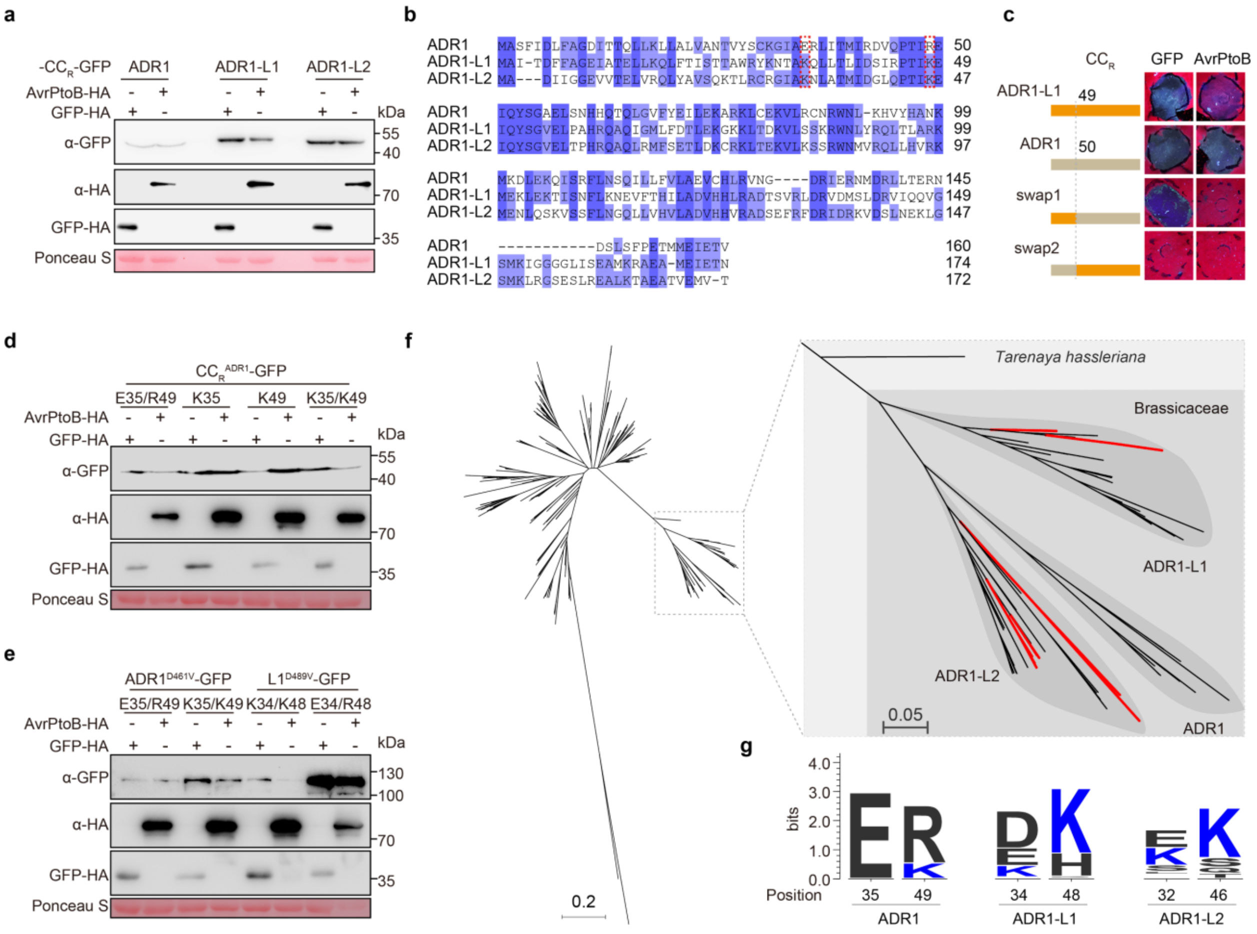


**Extended Data Fig. 3. Two lysine residues of the $CC_R$ domain are required for AvrPtoB-mediated suppression of ADR1-L1$^{D489V}$-triggered HR. a,** AvrPtoB induces the degradation of $CC_R^{ADR1-L1}$, but not $CC_R^{ADR1}$ and $CC_R^{ADR1-L2}$ in *N. benthamiana*. **b,** Alignment of $CC_R$ domains of the three ADR1 homologs. Dark blue, match in all three homologs; light blue, match in two homologs. The position of the ADR1-L1/ADR-L2-specific lysine residues K34 (K32) and K48 (K46) are marked by red boxes. **c,** The N-terminal 50 aa of $CC_R$ determine the specificity of AvrPtoB-mediated suppression of $CC_R^{ADR1-L1}$-triggered HR in *N. benthamiana*. **d,** AvrPtoB induces the degradation of $CC_R^{ADR1}$ (E35K/R49K) but not $CC_R^{ADR1}$ (E35K) and $CC_R^{ADR1}$ (R49K). **e,** AvrPtoB reduces the accumulation of full-length ADR1$^{D461V}$ (E35K/R49K) but not ADR1-L1$^{D489V}$ (K34E/K48R) in *N. benthamiana*. **f,** Maximum-likelihood phylogeny of 552 ADR1 homologs from angiosperms. ADR1 homologs from Brassicaceae fall into three clades, reflecting the three homologs found in Arabidopsis. The consensus tree was constructed using PhyML, with an NRG1-like protein as outgroup. ADR1-L1 and ADR1-L2 homologs with lysines at both sites are highlighted in red. **g,** Frequency of the two key residues of $CC_R$ required for AvrPtoB targeting among different ADR1 homologs from Brassicaceae.

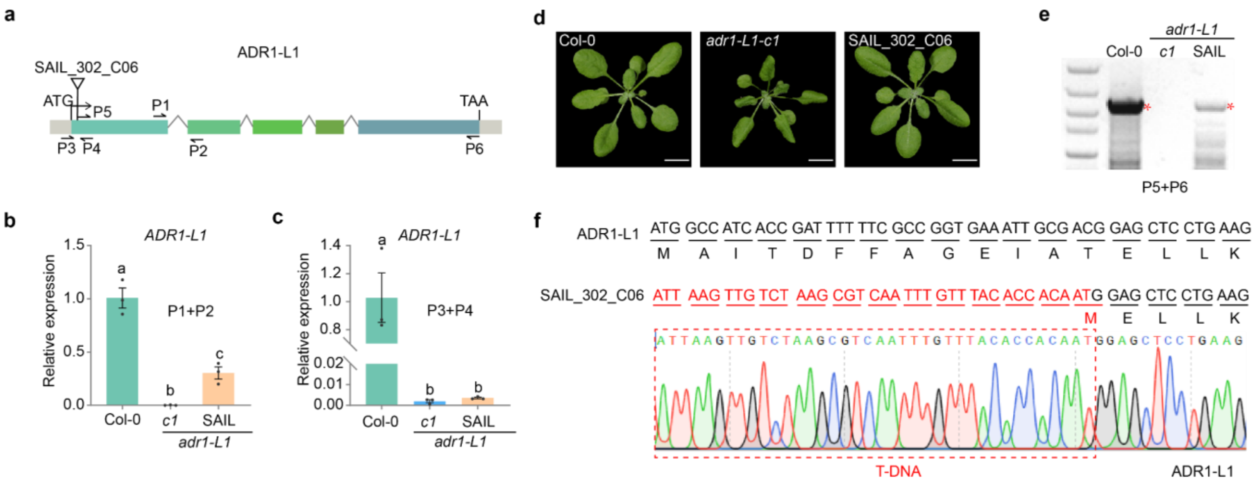


**Extended Data Fig. 4. Differences in morphology between the *ADR1-L1* T-DNA insertion line *SAIL_302_C06* and *acr1-L1-c1* grown under the same conditions.** **a,** Diagram of primers for genotyping for T-DNA insertion and *ADR1-L1* RT-PCR. The primer pair P1+P2 was used for measuring RNA expression from an *ADR1-L1* fragment outside of the T-DNA insertion. The primer pair P3+P4 was used for measuring RNA from an *ADR1-L1* fragment across the T-DNA insertion. The primer pair P5+P6 was used for amplifying coding sequences transcribed from the *SAIL_302_C06* line. **b,** RNA expression from an *ADR1-L1* fragment outside of T-DNA insertion in *SAIL_302_C06* line. Data represent the mean and standard error of three biological replicates (n = 3 biologically independent samples, $p<0.05$, one-way ANOVA followed by Tukey's post hoc test, letters indicate significantly different groups). **c,** RNA expression of an *ADR1-L1* fragment across the T-DNA insertion in *SAIL_302_C06* line, as shown by qPCR experiment. Data represent the mean and standard error of three biological replicates (n = 3 biologically independent samples, $p<0.05$, one-way ANOVA followed by Tukey's post hoc test, letters indicate significantly different groups). **d,** The T-DNA insertion line SAIL_302_C06 line has normal morphology. Three-week-old Col-0, *adr1-L1-c1*, and *adr1-L1-1* SAIL_302_C06 plants grown at 23°C under short-day conditions are shown. Scale bar: 10 mm. **e,** Detection of expression of 5'-terminally truncated *ADR1-L1* coding sequences amplified by primers P5 and P6 in *SAIL_302_C06*, as shown by gel electrophoresis. **f,** Sequence of RT-PCR product from *SAIL_302_C06* confirms generation of an in-frame deletion of *ADR1-L1* coding sequences.

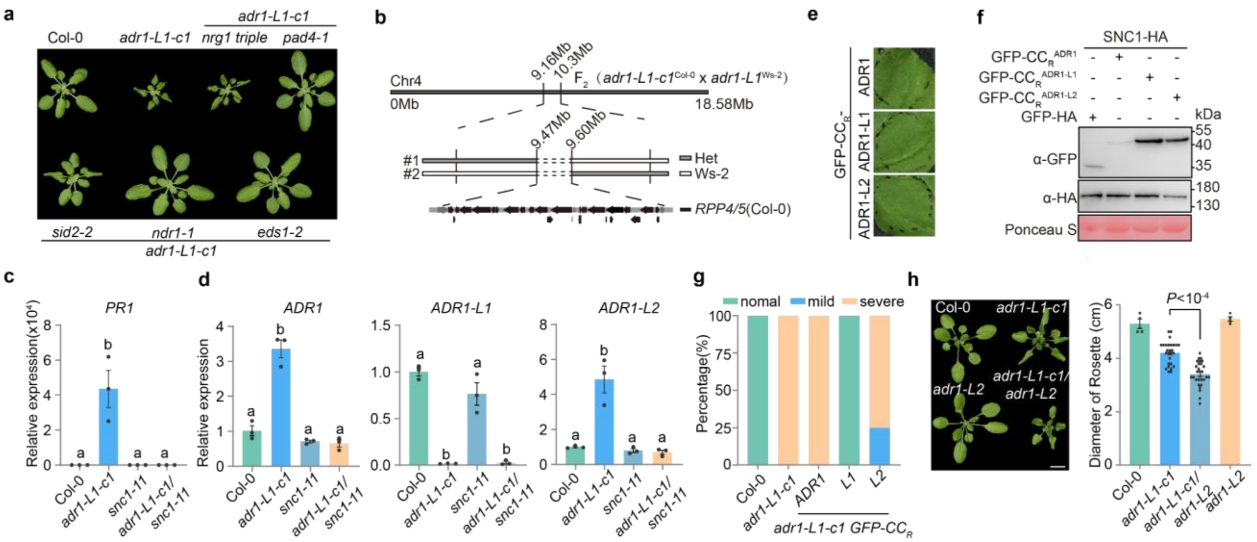


**Extended Data Fig. 5. *adr1-L1* null mutant defects are fully dependent on *SNC1*. a,** 3-week-old *adr1-L1-c1* in key ETI regulator mutant backgrounds, using Col-0 and *adr1-L1-c1* as the control, at 23°C under short-day conditions. Scale bar, 10 mm. **b,** Identification of *RPP4/5 NLR* cluster as the genetic locus responsible for suppression of *adr1-L1* defects in Ws-2, using an $F_2$ mapping population derived from crosses between *adr1-L1* (Ws-2) and *adr1-L1-c1* (Col-0). **c,** *PR1* expression in Col-0, *adr1-L1, snc1-11*, and *adr1-L1-c1/snc1-11*. **d,** Expression of *ADR1, ADR1-L1* and *ADR1-L2* in Col-0, *adr1-L1, snc1-L1*, and *adr1-L1-c1/snc1-11*. **e,** N-terminally GFP-tagged $CC_R$ domains of three ADR1 homologs do not induce cell death in *N. benthamiana*. **f,** Protein accumulation as shown by immunoblot for experiments shown in **Fig. 5e**. **g,** Distribution of different phenotypic classes in Arabidopsis $T_1$ plants transformed with *p35S::GFP-*$CC_R^{ADR1}$, *p35S::GFP-*$CC_R^{ADR1-L1}$, or *p35S::GFP-*$CC_R^{ADR1-L2}$ in the *adr1-L1-c1* background. **h,** An *adr1-L2* mutation enhances the autoimmune phenotype of *adr1-L1-c1*. Shown are3-week-old Col-0, *adr1-L1-c1*, *adr1-L2*, and *adr1-L1-c1/adr1-L2* plants grown at 23°C under short-day conditions (left panel), and their rosette diameters (right panel). Data represents the mean and standard error, which were analysed by two-tailed Student's t-test. Scale bar, 10 mm.
